# LASSI: A lattice model for simulating phase transitions of multivalent proteins

**DOI:** 10.1101/611095

**Authors:** Jeong-Mo Choi, Furqan Dar, Rohit V. Pappu

## Abstract

Biomolecular condensates form via phase transitions that combine phase separation or demixing and networking of key protein and RNA molecules. Proteins that drive condensate formation are either linear or branched multivalent proteins where multivalence refers to the presence of multiple protein-protein or protein-nucleic acid interaction domains or motifs within a protein. Recent work has shown that multivalent protein drivers of phase transitions are in fact biological instantiations of *associative polymers*. Such systems can be characterized by *stickers-and-spacers* architectures where stickers contribute to system-specific spatial hierarchies of directional interactions and spacers control the concentration-dependent inhomogeneities in densities of stickers around one another. The collective effects of interactions among stickers and spacers lead to the emergence of dense droplet phases wherein the stickers form percolated networks of polymers. To enable the calculation of system-specific phase diagrams of multivalent proteins, we have developed LASSI (**LA**ttice simulations of **S**ticker and **S**pacer **I**nteractions), which is an efficient open source computational engine for lattice-based polymer simulations built on the stickers and spacers framework. In LASSI, a specific multivalent protein architecture is mapped into a set of beads on the 3-dimensional lattice space with proper coarse-graining, and specific sticker-sticker interactions are modeled as pairwise anisotropic interactions. For efficient and broad search of the conformational ensemble, LASSI uses Monte Carlo methods, and we optimized the move set so that LASSI can handle both dilute and dense systems. Also, we developed quantitative measures to extract phase boundaries from LASSI simulations, using known and hidden collective parameters. We demonstrate the application of LASSI to two known archetypes of linear and branched multivalent proteins. The simulations recapitulate observations from experiments and importantly, they generate novel quantitative insights that augment what can be gleaned from experiments alone. We conclude with a discussion of the advantages of lattice-based approaches such as LASSI and highlight the types of systems across which this engine can be deployed, either to make predictions or to enable the design of novel condensates.

**Author Summary:** Spatial and temporal organization of molecular matter is a defining hallmark of cellular ultrastructure and recent attention has focused intensely on organization afforded by membraneless organelles, which are referred to as biomolecular condensates. These condensates form via phase transitions that combine phase separation and networking of condensate-specific protein and nucleic acid molecules. Several questions remain unanswered regarding the driving forces for condensate formation encoded in the architectures of multivalent proteins, the molecular determinants of material properties of condensates, and the determinants of compositional specificity of condensates. Building on recently recognized analogies between associative polymers and multivalent proteins, we have developed and deployed LASSI, an open source computational engine that enables the calculation of system-specific phase diagrams for multivalent proteins. LASSI relies on *a priori* identification of stickers and spacers within a multivalent protein and mapping the stickers onto a 3-dimensional lattice. A Monte Carlo engine that incorporates a suite of novel and established move sets enables simulations that track density inhomogeneities and changes to the extent of networking among stickers as a function of protein concentration and interaction strengths. Calculation of distribution functions and other nonconserved order parameters allow us to compute full phase diagrams for multivalent proteins modeled using a stickers-and-spacers representation on simple cubic lattices. These predictions are shown to be system-specific and allow us to rationalize experimental observations while also enabling the design of systems with bespoke phase behavior. LASSI can be deployed to study the phase behavior of multicomponent systems, which allows us to make direct contact with cellular biomolecular condensates that are in fact multicomponent systems.

## Introduction

Biomolecular condensates organize cellular matter into distinct non-stoichiometric assemblies comprising of proteins and nucleic acids (1). Prominent condensates include nuclear bodies (2) such as the nucleolus, nuclear speckles (3, 4), and germline granules (1, 5, 6). Condensates also form in the cytoplasm. These include stress granules (7), membrane-anchored signaling clusters (8, 9), and bodies in post-synaptic zones (10). All of these condensates share key features: (i) they range in size from a few hundred nanometers to tens of microns (1, 2, 11); (ii) they are multicomponent entities comprising of hundreds of distinct types of proteins and nucleic acids; (iii) and of the hundreds of different types of molecules that make up condensates, a small number perform the role of *scaffolds* that seem to be essential for the formation of condensates, while the remainder are *clients* that partition selectively into condensates once they are formed (1, 12). The simplest feature that distinguishes protein scaffolds from clients is the valence of interacting domains / motifs. Scaffolds have higher valence of interaction domains / motifs when compared to clients (1, 12–14).

Biomolecular condensates form and dissolve in an all-or-none manner (2, 11, 15). The reversible formation and dissolution of condensates can be controlled by the concentrations of scaffold molecules; condensates form when concentrations of scaffold molecules cross scaffold-specific threshold values known as *saturation concentrations* (15). The transitions that characterize condensate formation bear the hallmarks of a sharp transition in scaffold density, leading to the formation of a dense phase that is in equilibrium with a dilute phase. This type of transition, known as *phase separation*, sets up two or more coexisting phases to equalize the dense and dilute phase chemical potentials of scaffold molecules across phase boundaries (15). Phase separation is reversible and this reversibility can be achieved by (i) changes to concentrations of scaffold molecules (9, 16), (ii) changes to solution conditions that alter the effective interaction strengths among scaffold molecules (17–20), (iii) altering saturation concentrations through ligand binding – a phenomenon known as polyphasic linkage (21, 22), or (iv) via biological regulation such as post-translational modifications of proteins (8, 12, 23).

Recent studies have focused on uncovering the defining features of scaffold proteins (13, 15, 17–19, 24–40) and RNA molecules (41–43). These studies suggest that *scaffolds* are biological instantiations of *associative polymers* (44) characterized by a so-called *stickers-and-spacers* architecture (45). Stickers contribute to a hierarchy of specific pairwise and higher-order interactions whereas spacers control the concentration-dependent inhomogeneities in the densities of stickers around one another. Stickers can be *hot spots* or *sectors* (46) on the surfaces of folded proteins (15, 29) or short linear motifs within intrinsically disordered regions (15, 24, 47). Spacers are typically intrinsically disordered regions that contribute through their sequence-specific effective solvation volumes to the interplay between *density transitions* (phase separation) and *networking transitions* (percolation or gelation) (28, 29). Spacers can also be folded domains that are akin to uniformly reactive colloidal particles, although this has not been explored in any kind of detail. Protein scaffolds map to stickers-and-spacers architecture as linear multivalent proteins, branched multivalent proteins, or some combination of the two (13, 15).

A typical two-component system comprises of the solvent (which includes all components of the aqueous milieu) and a scaffold molecule. For fixed solution conditions, one can generate phase diagrams (25) as a function of protein concentration, the valence of stickers, the affinities of stickers, the sequence-specific effective solvation volumes of spacers, and the lengths / stiffness of spacers. The resultant five-dimensional phase diagram can be projected onto a plane whereby one investigates the phase behavior while keeping the valence of stickers, the lengths of spacers, and effective solvation volumes of spacers fixed while varying the concentration of stickers and the affinities between stickers (29). Changes to protein concentration will enable density fluctuations and above the saturation concentration, designated as *c*_sat_, the density inhomogeneities lead to separation of the system into coexisting phases. The concentration of multivalent proteins in the dilute and dense phases will be denoted as *c*_sat_ and *c*_dense_, respectively. For a given bulk concentration *c*_bulk_ between *c*_sat_ and *c*_dense_, the fraction of molecules within each of the coexisting phases is governed by the lever rule (48).

Stickers also form reversible physical crosslinks and these crosslinks generate networks of inter-connected proteins. The number of proteins within the largest network of the system grows continuously as the protein concentration increases. Above a concentration threshold known as the percolation threshold and designated as *c*_perc_, the single largest network spans the entire system and this phenomenon is called *percolation* (49–51). If the percolated networks have the rheological properties of viscous liquids or viscoelastic fluids, the fluids are referred to as *network fluids* (15, 52).

Phase separation and percolation can be coupled to one another. The coupling will depend on the values of *c*_sat_, *c*_dense_, and *c*_perc_ relative to *c*_bulk_. If *c*_bulk_ is smaller than all of *c*_sat_, *c*_dense_, and *c*_perc_, the system is in a single dilute phase with no large molecular networks (**Fig 1a**). If *c*_bulk_ > *c*_perc_ and *c*_perc_ < *c*_sat_, then a system-spanning percolated network forms without phase separation (**Fig 1b**). However, the system undergoes phase separation and a dense phase forms as a *percolated droplet* if *c*_bulk_ > (*c*_sat_, *c*_perc_) and *c*_sat_ < *c*_perc_ < *c*_dense_ (**Fig 1c**). Recent studies, using three-dimensional lattice models designed to mimic the poly-SH3 and poly-PRM systems of Li *et al*. (16), show that sequence-specific effective solvation volumes of linkers / spacers between folded domains directly determine whether phase separation and percolation are coupled or if percolation occurs without phase separation for linear multivalent proteins (28, 29). The coupling between phase separation and percolation is controlled by the extent to which spacers / linkers preferentially interact with the surrounding solvent.

**Fig 1.**
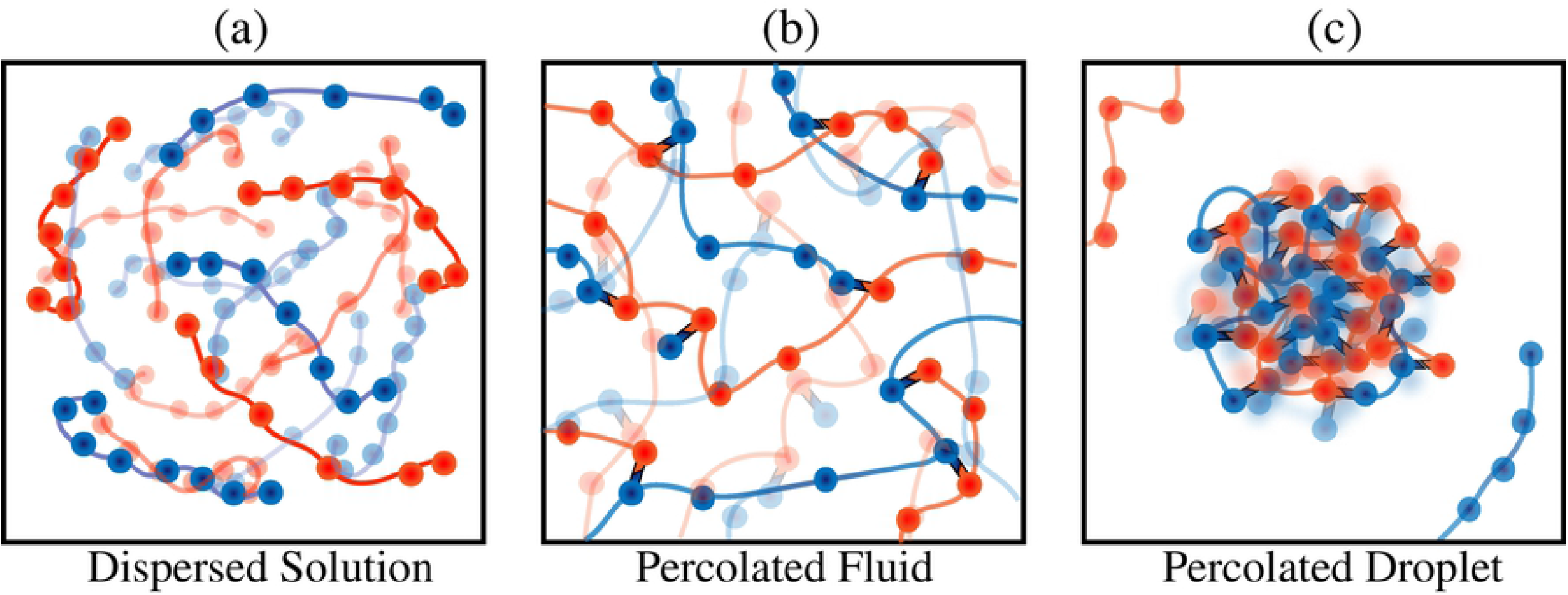
Characteristic phases in the stickers and spacers formalism. (a) Dispersed solution phase where the polymers are uniformly mixed in solution. (b) Percolated fluid wherein the polymer chains form a percolated, system-spanning network through physical crosslinks among stickers results. (c) Droplet wherein the network formation also causes the polymers to collapse onto each other.

Lattice models have also been adapted to model the phase transitions for other systems comprising of different numbers of multivalent protein and RNA molecules (28–30, 43, 53–55). In the present work, we provide a formal description of the design and implementation of system-specific lattice models for simulating phase transitions of multivalent proteins. The simulation engine, known as LASSI for **LA**ttice simulations of **S**ticker and **S**pacer **I**nteractions, formalizes the approaches that have been developed and deployed in recent studies (28–30, 53, 54). We describe the design of LASSI, focusing first on the overall structure of the model, the Monte Carlo sampling, and their justification for generic multivalent proteins. We further describe the calculation of order parameters for quantifying phase separation and percolation. Then, using two specific examples of linear and branched multivalent protein systems, we illustrate the deployment of LASSI to two biologically relevant systems. In both systems, we make *a priori* assumptions regarding the identities of stickers and spacers, which is a requirement for the deployment of LASSI. Although we focus here primarily on systems with a few components, it should be emphasized that the design of LASSI is able to handle a range of multicomponent systems.

## Materials and Methods

Considerations that go into the development of a suitable lattice model include (a) the choice of the *mapping* between a specific multivalent protein of interest and a lattice representation, (b) the *parameterization* of the strengths and ranges of interactions for all unique pairs of beads and vacancies, (c) the *design* of move sets and acceptance criteria for Monte Carlo simulations that enable the sampling of local and collective motions of large numbers of lattice-instantiated multivalent proteins, (d) the efficient *titration* of key parameters such as protein concentrations and interaction strengths, and (e) the *extraction* of phase boundaries in terms of known and hidden collective parameters, which become the relevant *order parameters* for phase transitions of interest.

### Generating lattice representations of multivalent proteins

For a given linear or branched multivalent protein, we first choose a suitable mapping between the protein degrees of freedom and a lattice representation. The conformational space is a simple cubic lattice with periodic boundary conditions used to mimic a macroscopic system. Phase transitions represent the collective effects of large numbers of molecules, and simulations have to include at least 10^3^ – 10^4^ protein molecules to observe facsimiles of these collective transitions in finite sized systems (56). Further, we need to be able to test for the effects of finite size artifacts and this requires a titration of the effects of varying the numbers of molecules. In effect, the lattice has to be large enough to accommodate at least 10^3^ molecules of each type for the most dilute concentrations. Often, we might need to increase the number of molecules to be of *O*(10^4^). Accordingly, a one-to-one mapping between the protein degrees of freedom and a lattice representation would lead to a computationally intractable model. Instead, we adopt system-specific coarse-graining approaches, whereby the coarse-graining is guided by *a priori* rigorous or phenomenological knowledge of the identities of *stickers* versus *spacers*. For disordered proteins, the stickers within disordered regions often correspond to single amino acid residues or short linear motifs. For multivalent folded proteins, the stickers are either an entire protein domain or sectors on domain surfaces (28, 29). Residues corresponding to spacers may either be modeled explicitly, where one or more spacer residues are modeled by a single bead on the lattice site, or be modeled as phantom tethers, where the intrinsic lengths of tethers are calibrated in terms of the numbers of lattice sites (28, 29). In both cases, the tethers can stretch, bend and flex and these degrees of freedom contribute to density inhomogeneities that are the result of altered patterns of inter-sticker interactions.

### LASSI and Bond Fluctuation Models

The structure of LASSI is inspired by the bond fluctuation model (BFM) for lattice polymers (57). This is a general lattice model for simulations designed to extract equilibrium conformational distributions and dynamical attributes of polymers in dilute solutions as well as dense melts. There are two versions of the BFM viz., the Carmesin-Kremer BFM or CK-BFM (58) and the Shaffer BFM or S-BFM (57). Both models are based on the use of simple cubic lattices, which discretizes the conformational space for polymers.

In the CK-BFM (58), each repeating unit or monomer within a polymer is modeled as a cube that occupies eight 3-dimensional lattice positions and bond vectors connect pairs of monomers. Overlap of monomers is associated with an energetic penalty, and each bond vector can have up to 108 distinct directions. The choice of bond vector set encodes the geometry of the polymer and places constraints on the bond lengths and bond angles. All other interactions are governed by the inter-monomer potentials, and evolution of the system through conformational space is driven by the changes to the overall potential energy. In contrast, the S-BFM places each monomer on a single lattice site. Covalently bonded monomers are connected by bonds that are constrained to be of three types, leading to chains that have bonds of length 1, 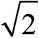 or 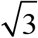 in units of lattice size. Monte Carlo moves with suitable acceptance criteria can be designed for both types of BFMs. The simulations are used to generate equilibrium conformational distributions of lattice polymers in either dilute or dense phases. The move sets control the overall polymer dynamics and the acceptance of different types of moves and the calculation of correlation functions allows one to compute dynamical quantities for lattice polymers (57). If we were to use either of the established BFMs without modification, then each amino acid residue would be modeled as a monomer, and such an approach would be useful when the identities of stickers and spacers remain ambiguous and this approach underlies a different simulation engine known as PIMMS (43).

LASSI is a generalization of the S-BFM that also adapts features of the CK-BFM. Given a choice of the mapping for the coarse-graining procedure, each multivalent protein is described as a chain of non-overlapping monomers viz., beads that occupy sites on a 3-dimensional cubic lattice. Note that the choice of a single site per bead is similar to that of the S-BFM, although the bead, which is a sticker or spacer monomer, need not be the monomeric unit, *i.e*., an amino acid residue in the case of proteins. Each sticker monomer is linked to its adjacent sticker on the chain via either a phantom tether or a set of spacer beads that occupy individual lattice sites (28, 29). A spacer / tether length of unity implies that adjacent monomers are within 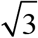 lattice units of one another (**Fig 2**). The choice of the spacer length will be sequence-specific.

**Fig 2.**
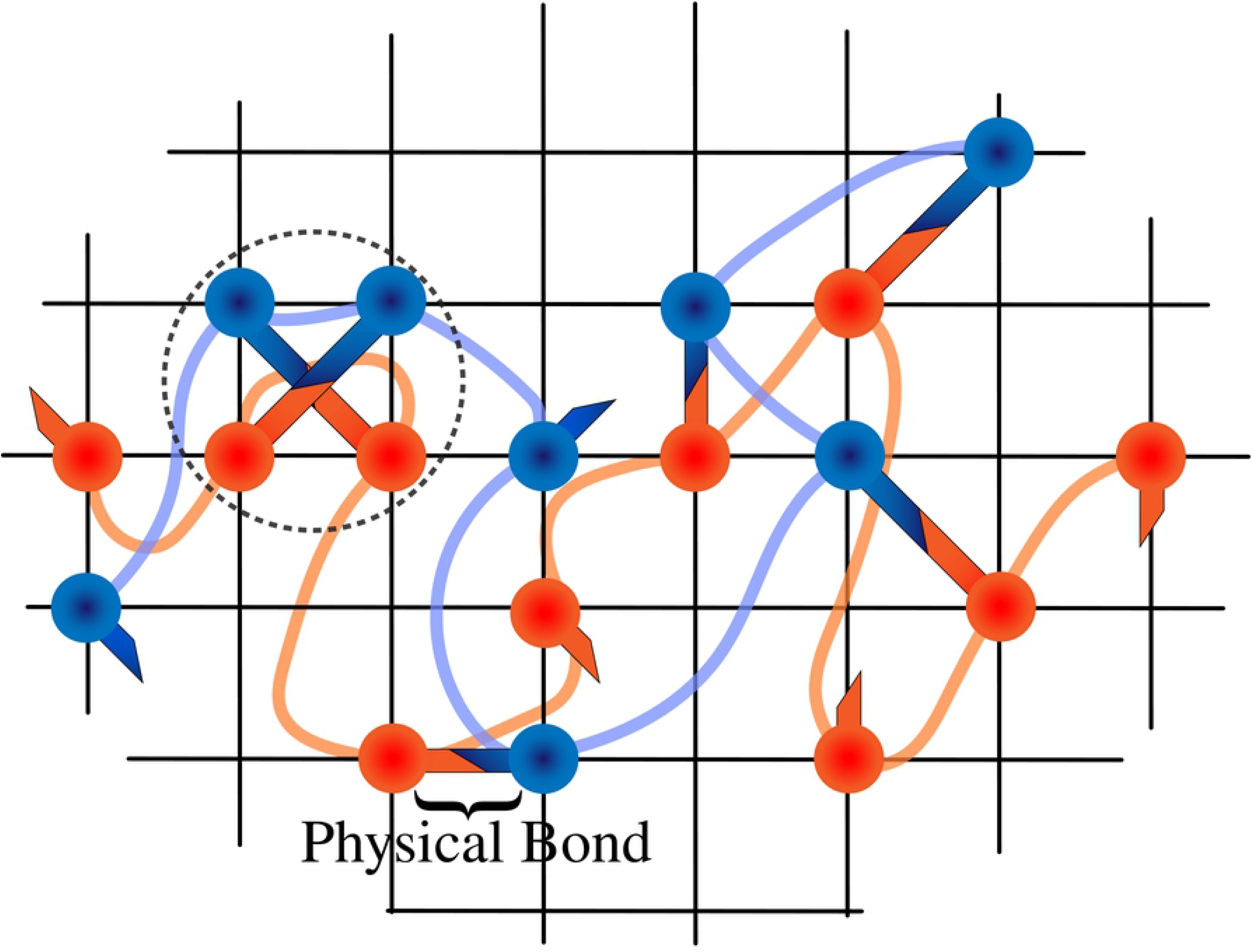
2-dimensional representation of the LASSI architecture. The beads with arms denote stickers where arms denote that the monomers are capable of orientational interactions, and the curved lines connecting the monomers represent phantom tethers, which are allowed to freely overlap (implicit spacer model). Different colors denote different sticker and spacer species respectively. Note that the physical bonds are allowed to overlap (dashed circle). For the rest of this paper, physical bonds will not be labeled and will only be depicted as overlapping orientational arms.

Inter-monomer (sticker-sticker, sticker-spacer, and spacer-spacer) interactions are modeled as contact-based pairwise interactions. A sticker monomer can bind to another sticker monomer that occupies an adjacent lattice site with an interaction energy that depends on the types of both monomers. Monomers are considered to be adjacent to one another if they are within a lattice distance of 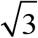. By this criterion, each lattice site occupied by a sticker monomer will have 26 adjacent lattice sites. This is reminiscent of the interaction geometry of a CK-BFM for each monomer. In the current implementation of LASSI, the interactions are mutually exclusive, implying that a sticker cannot interact simultaneously with more than one other sticker, even though there are 26 adjacent sites that the interaction partner can occupy. The unoccupied sites for each sticker will effectively contribute to an overlap cost, if the sticker is already engaged in another inter-monomer interaction with stickers or spacers. The combination of the geometry of the interaction sites per monomer and the single occupancy constraint leads to anisotropic interactions between sticker interactions. This feature is unique to LASSI and it is not incorporated in other variants of BFMs and allows us to deploy LASSI for modeling heteropolymeric systems.

### Setup of simulations

A system with *n* multivalent proteins is in reality an *n*+1 component system since the solvent is the hidden or implicit component. In LASSI, sites that are not occupied by protein units automatically represent solvent sites. Although the interaction potentials do not explicitly include terms between solvent and protein sites, the effective interaction strengths between pairs of protein units represent an averaging over protein-protein, protein-solvent, and solvent-solvent interactions. The solvent sites, *i.e*., the sites that are not occupied by protein units, represent contributions from the solvent to the overall translational and mixing entropies. Simulations are initiated by randomizing the positions of protein units, subject to the constraints of chain connectivity.

The parameters that are set at the start of each LASSI simulation include the total number of molecules *n_i_* of type *i* and the size of the lattice *L*, from which we can calculate the total number *n* of all protein components 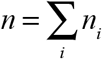 and the concentration or number density of each protein *c_i_* = *n_i_*/*L*^3^. The setup also includes stipulations for the architectures of each of the protein such as specification of the number of monomers per chain, the overall topology of each protein (linear *vs*. branched), the lengths of spacers, and the types of spacers (implicit / phantom vs. explicit) (28–30, 53, 54). The number of monomers will equal the sum of the number of stickers and spacers if spacer residues are modeled explicitly. Alternatively, if spacers are modeled as phantom tethers, then the number of explicitly modeled monomers will equal the number of stickers. Specification of the energetics of the system includes specification of the simulation temperature in normalized units, homotypic and heterotypic interaction strengths between pairs of stickers, the energetic cost for the overlap of stickers, and the interaction strengths between sticker and spacer sites if the spacers are modeled explicitly.

### Design of Monte Carlo move sets

The design objective of LASSI is to compute sequence- and architecture-specific phase diagrams for systems comprising of one or more types of linear or branched multivalent proteins. This requires a simulation strategy that enables the sampling of the full spectrum of coexisting densities and networked states for multivalent proteins. Accordingly, the conformations of randomly initialized systems of proteins on a simple cubic lattice are sampled via a series of Markov Chain Monte Carlo (MCMC) moves that are designed to ensure efficient sampling of changes in protein density and networking while maintaining microscopic reversibility. We developed and deployed a collection of moves and these are described below.

#### Monte Carlo sampling with biases

In LASSI, we have independent contributions from two main energetic sources. Monomer units are not allowed to overlap, and this can be described by a position-dependent energy *E*_pos_ where *E*_pos_ = 0 or ∞. On the other hand, inter-monomer pairwise interactions also contribute to the total energy, and *E*_rot_ denotes the sum over all of the effective pairwise inter-monomer interaction energies. The subscript “rot”(rotational) indicates the fact that for a pair of nearest neighbor stickers their interaction energies are actually governed by their mutual orientations. Accordingly, the total system energy in a specific configuration *i* is written as:

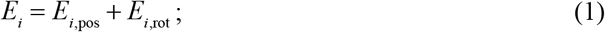

The equilibrium probability associated with configuration *i* is given by the Boltzmann distribution as:

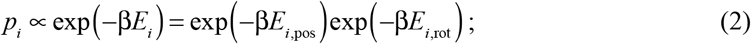

In equation (2), β is the inverse of the simulation temperature in units of the Boltzmann constant (effectively, *k_B_* = 1 energy unit / temperature unit). The frequency with which a transition from configuration *i* to *j* is proposed will be governed by the elements *T_ij_* of the targeting matrix **T**. The proposed transition is accepted / rejected based on the elements *A_ij_* of the acceptance ratio matrix **A**. A MCMC move that transitions the system from configuration *i* to *j* defines a flow in configuration space and this flow has to satisfy detailed balance also referred to as microscopic reversibility:

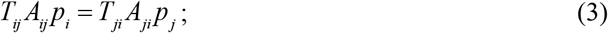

If the targeting matrix is symmetric, then *T_ij_* = *T_ji_* and the acceptance ratios such as those prescribed by Metropolis *et al*. (59) will ensure the preservation of detailed balance. However, if biases are incorporated into the targeting matrix, which is often essential to enhance the sampling of configurations that contribute to density inhomogeneities and the making / breaking of bonds in dense networks, then the elements of the acceptance ratio matrix have to be designed to ensure the preservation of detailed balance. We deploy a general strategy of using biased moves to enhance the sampling of different mutual orientations among pairs of stickers. The incorporation of these orientational biases is accounted for by modifying the acceptance criterion of Metropolis *et al*. (59) whereby each element of A is written as:

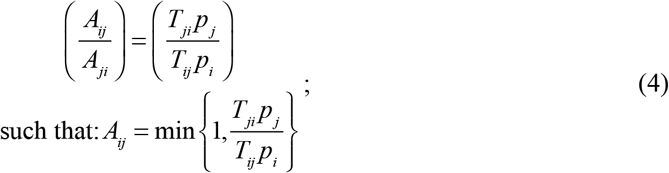

For a symmetric targeting matrix, we recover the standard acceptance ratio of Metropolis *et al*. (59) *viz*.,

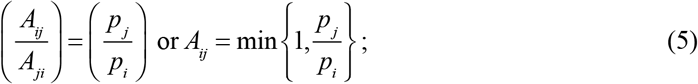

Since the moves within LASSI generally involve orientational biases, the elements *T_ij_* are rewritten in terms of a Rosenbluth weighting factor *W_j_* (60, 61) whereby:

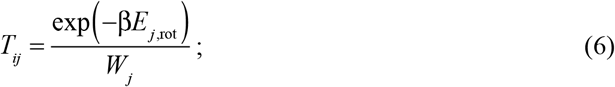

Substituting (6) into (4) leads to:

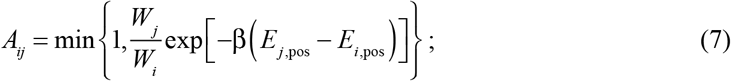

The specific form for the weighting factors *W_i_* will depend on the type of move because the extent of asymmetry in the targeting matrix will depend on the nature of the bias incorporated into the biasing move that proposes a transition from *i* to *j*. The specific forms for weighting factors are discussed in the context of the move types that are introduced next.

#### Rotational moves

A monomer is in an associated or a dissociated state and in the associated state it has a specified binding partner. This defines the rotational state of a monomer. To change the rotational state, we randomly pick a monomer from the system, and exhaustively sample all 26 adjacent sites to construct a list of potential binding partners. The rotational state of the monomer is changed, at random, by drawing a random integer *k* from a uniform distribution between [0, *b*], where *b* is the number possible binding partners available to the monomer. If *k* = the monomer is set to be in a dissociated state. Otherwise, the *k*^th^ candidate bond is formed and the state of the monomer is set to be in an associated state. If the monomer cannot be involved in a rotational interaction, as would be the case for an explicitly modeled spacer, the rotational move is rejected outright. The accessible volume for rotational interactions is within a cube of unit volume centered on the randomly chosen monomer (see **Fig 3** for a 2-dimensional representation), and hence, each sticker monomer will have *b*_max_ = 3^3^–1 possible sites as neighboring sites at most. Since this number is not large, we sample all 26 possible interaction sites.

**Fig 3.**
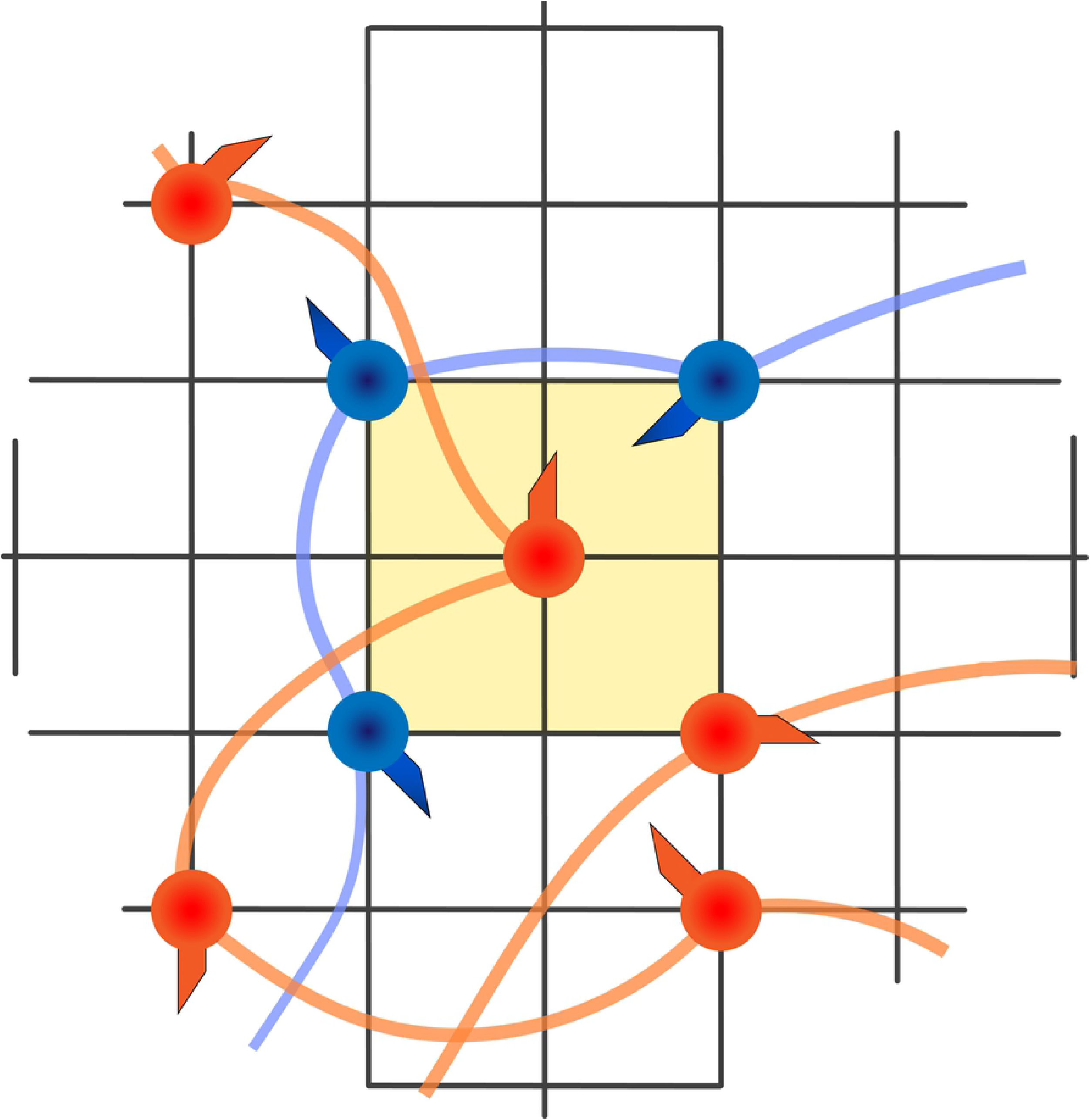
2-dimensional representation of rotation move. For a given randomly selected monomer (middle orange bead), 3^3^−1 nearest lattice sites (yellow box) are checked for possible interaction candidates, where eligible candidates have a non-zero interaction energy with the selected monomer. In this figure, orange stickers interact with blue stickers and thus this sticker has 3 possible candidates. The end orientational state of the monomer is then picked using the metropolis criterion, which also includes the non-interacting state.

#### Local moves

This move serves as the basic unit of local displacement of monomers – be they stickers or spacers. A randomly chosen monomer is moved from position **r**_*i*_ to **r**_*j*_ = **r**_*i*_ + Δ**r**. Acceptance of the move is predicated on the move not leading to an overlap with a site occupied by another monomer and the satisfaction of linker constraints. The choice for Δ**r** is made by uniformly sampling each component from the interval [−2,2] such that 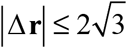, as shown in **Fig 4**. Note that these local moves are analogous to *crankshaft* or *kink jump* moves if the selected monomer is in the interior of a molecule (62), and they become analogous to *end rotation* moves if end monomers are selected (63).

**Fig 4.**
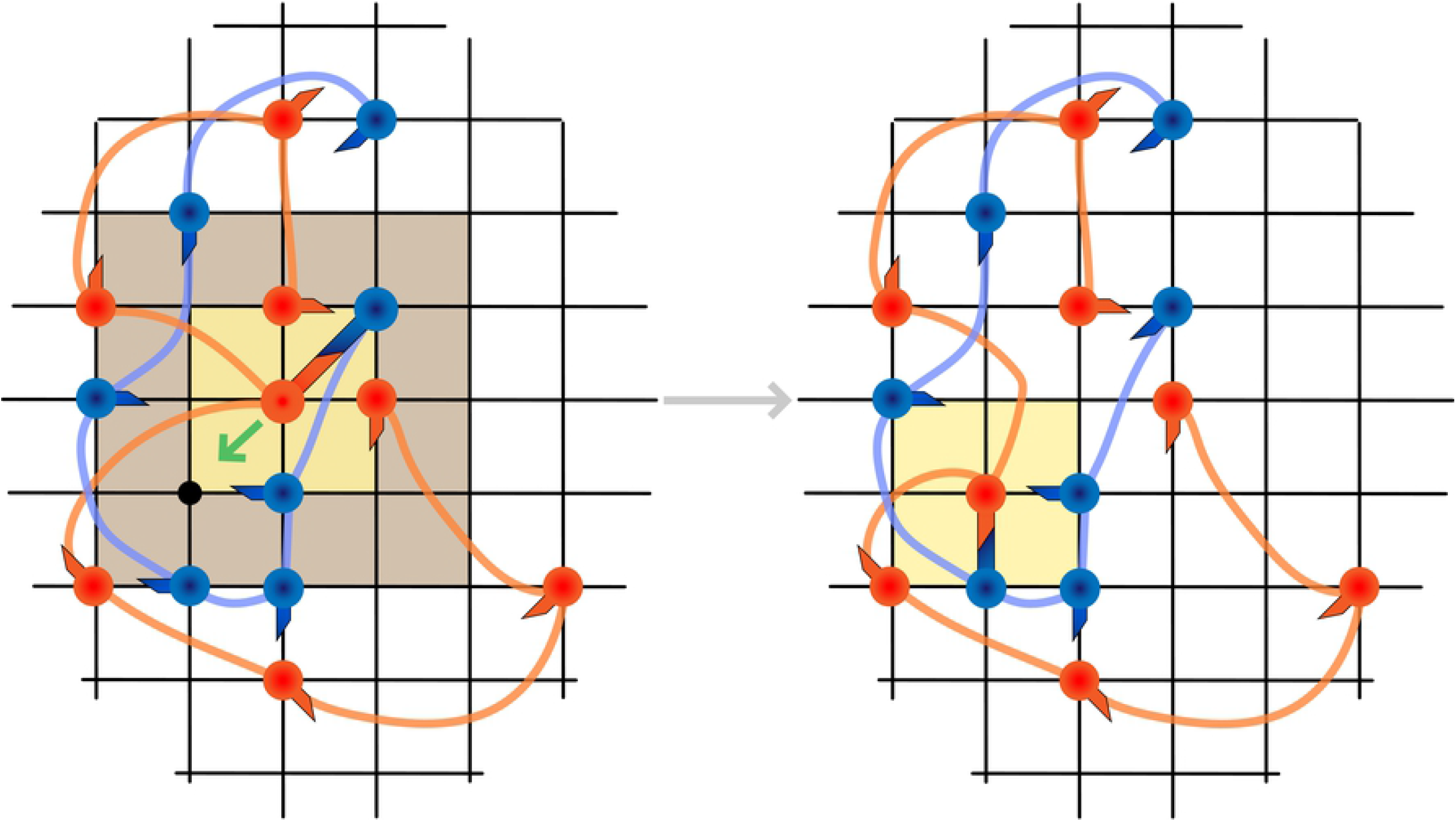
2-dimensional representation of local move. For a given randomly selected monomer, a new location is proposed by sampling ±2 lattice sites in each coordinate (brown box) and picking a lattice site that is empty. If an empty lattice site is found within a pre-determined number of trials, the numbers of interacting candidates are calculated at the old and proposed location (yellow boxes). Then the move is accepted or rejected using the modified Metropolis criterion that considers orientational bias (see text).

Local moves have a rotational bias in LASSI and the Rosenbluth factor is calculated as follows. Starting with equation (7), we shall designate the chosen monomer have index *k*. In configuration *i*, assume that monomer *k* has a binding partner of index *l*. The energy associated with the bond between monomers *k* and *l* is written as ε_*t*(*k*)*t*(*l*)_, where *t*(*x*) indicates the type of monomer *x*. The local move causes a change in binding partner, whereby the monomer *k* now binds to monomer *m*. The local move leads to a bond swap that causes a change in rotational energy, which is written as:

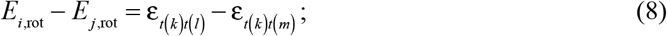

Use of equation (8) in equation (6) leads to:

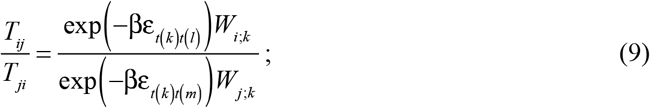

In equation (9), each Rosenbluth weight has an additional index in the subscript to indicate that the change in configuration is achieved by a change in the binding partner for the monomer *k*. To accelerate the creation of density inhomogeneities in supersaturated systems and facilitate the making and breaking of networks, we decompose *W_i;k_* as:

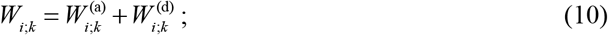

The two terms in equation (10) respectively represent the contributions to the Rosenbluth weights for monomers in associated (a) and dissociated states (d). First, we calculate the weight factors for the interacting monomers as a partition function over all nearest neighbor contributions, such that:

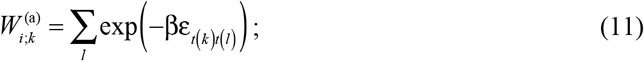

In equation (11), the summation runs over all potential binding partners *l* (nearest neighbors) for the monomer *k*. To illustrate how the Rosenbluth factor is calculated, we assume that the system has only one type of interactions with the pairwise energy designated as ε. If the number of nearest neighbors for monomer *k* in configuration *i* is designated as *N_k;i_*, then the Rosenbluth weight factor in equation (11) becomes:

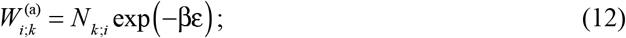

Setting 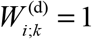 to incorporate a bias towards associated states, and using the simplification that leads to equation (12), we rewrite equation (9) as a definition of the acceptance criterion for the move from configuration *i* to *j* via local move involving monomer *k* as:

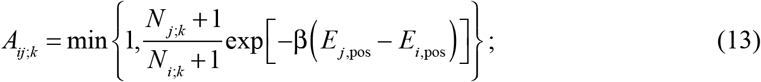

This choice for the acceptance criterion ensures that detailed balance is preserved while enhancing the sampling of configurations characterized by the breaking of old bonds and the forming of new ones.

#### Reptation – or slithering snake – moves

In dense configurations, it becomes difficult to realize large-scale translational or rotational motions of polymers. The *slithering snake move* is a Monte Carlo instantiation of reptation as first conceived by de Gennes (64). In this move, a chain is chosen at random, and the monomer at one end of the target chain is moved to a new position. The remaining monomers within the target chain are then successively moved such that monomer *m* along the chain moves into the previous position of monomer *m*–1 (**Fig 5**). This move relies on an inherent symmetry of chain molecules, because bond lengths between monomers are the same; if one swaps monomers across chains, the identity of the chain remains invariant. However, this move cannot be used if the molecule has heterogeneous bond lengths or if it is a branched polymer.

**Fig 5.**
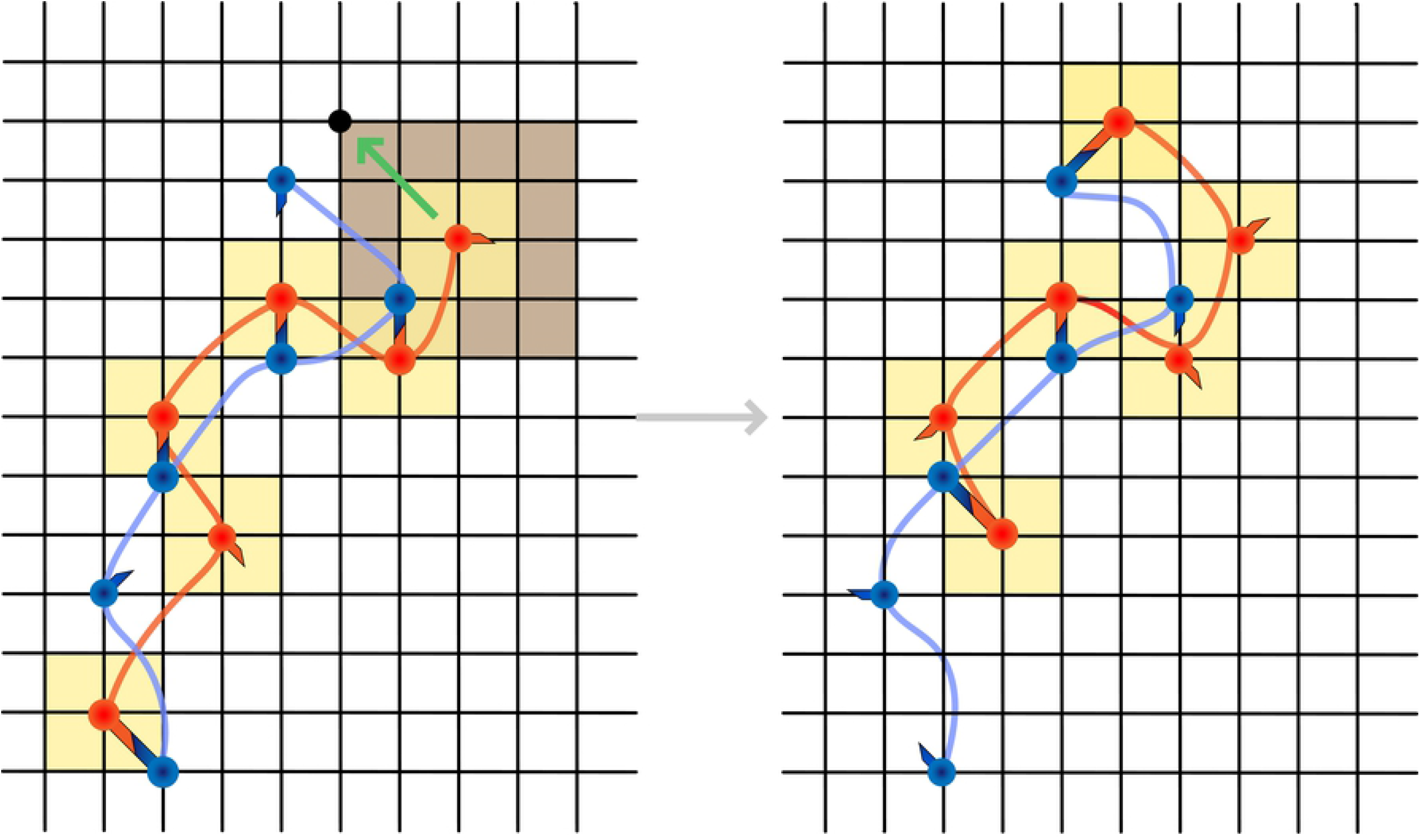
2-dimensional representation of reptation move. For a given randomly selected chain that has the same linker lengths between each monomer, an end is randomly picked. Then, a version of the local move is performed where the selected end is moved to a new random location that is an empty lattice site within 2 lattice sites in each coordinate (brown box). If an empty site is found within a predetermined number of trials, the number of orientational candidates is calculated for the whole chain in the old and the new configuration (yellow boxes). The modified metropolis criterion is then used to determine if the move is accepted or rejected. Note that since the whole chain is orientationally biased, monomers may have a different orientational state after the move is accepted, as shown in the figure.

The reptation move is rotationally biased, and this is true for every monomer in a chain. The bias is independent for each monomer and accordingly, the Rosenbluth factor for a single reptation move can be calculated from the Rosenbluth factors for each monomer-specific local move. In configuration *i* we obtain:

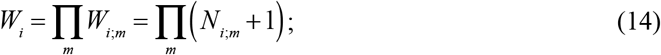

In equation (14), the product runs over all monomers *m* within the chain of interest. The acceptance criterion for a reptation move takes the form:

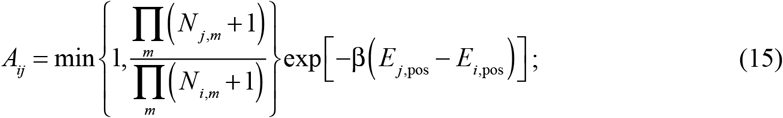

The inclusion of the bias for every interacting monomer, rather than just the end monomers, is to emulate how a real transiently bonded polymer would slither along its contours. Note that for strict detailed balance, the Rosenbluth factors for the two end monomers should be calculated and added to the acceptance criterion, but the current implementation of LASSI uses the first trial position that satisfies the position constraints.

#### Double pivot moves

These moves swap a part of a chain with the corresponding part of another chain of the same type. A monomer is picked at random; it is denoted as *i_m_*, where *i* is the monomer index within a chain, and *m* is the chain index. A search is then performed around the monomer within a prescribed distance for a monomer within the same type of chain. The requirement for the search is that the monomer of interest be one index ahead along its own chain, (*i*+1)_*n*_. Next a check is performed to ensure that the distance between *i_m_* and (*i*+1)_*n*_ is within the bond constraint connecting *i_m_* to (*i*+1)_*m*_, and that the distance between (*i*+1)_*m*_ and *i_n_* is within the bond constraint for *i_m_* and (*i*+1)_*m*_. Each monomer (*i*+1)_*n*_ that satisfies each of these constraints is stored and one of these is randomly picked for the double pivot move (**Fig 6**). The move is always accepted if there is a candidate because only connectivity changes unless bonds are modeled using elastic springs.

**Fig 6.**
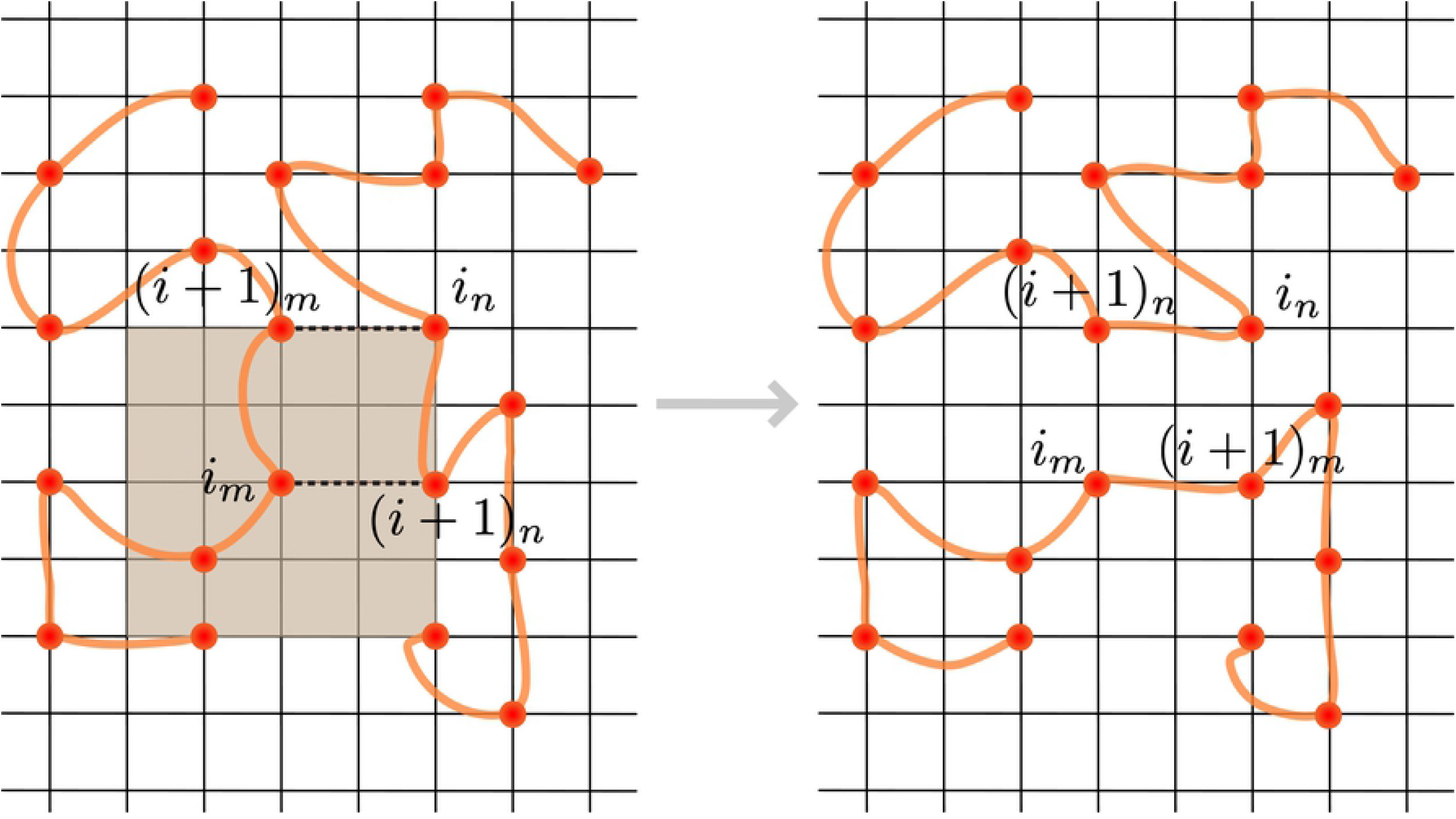
2-dimensional representation of double pivot move. For a randomly selected monomer, a 2×2×2 cube around the monomer is searched for appropriate bridging candidates (brown box), where an appropriate bridging candidate is the next monomer from a different chain, is within a linker length of the selected monomer as shown by the dashed line connecting *i_m_* and (*i*+1)_*n*_. Furthermore, the distance between (*i*+1)_*m*_ and *i_n_* must also be within a linker length as depicted by the upper dashed line. A list of all possible candidates is calculated and then a randomly chosen candidate is used to break and remake covalent bonds. This results in a large conformational change for both polymers. If the selected polymer is not linear, the move is rejected outright.

The purpose of this move is to engender large configurational changes in dense polymer melts, which approximate the dense phases formed upon phase separation. In dense regions the rate of acceptance of local moves decrease precipitously. At high enough densities, polymers become entangled and local moves reduce to slithering-snake moves and polymers are restricted to motions along tubes around one another (65, 66). Therefore, rather than physically moving polymers to create a change in configurations, we incorporate move sets that break and recreate bonds while ensuring that monomers do not overlap, and that bond constraints are always satisfied. If two chains are close enough to each other that the bonds between two monomers can be swapped, then such the double pivot move results in a large configurational change for both chains, and for the system.

#### Chain and cluster translation moves

The chain translation move is designed to move single chains while forming new bonds at the proposed location. This move attempts to translate the center of mass of a chain *i* from **r**_*j*_ to **r**_*j*_ = **r**_*i*_ + Δ**r** where 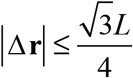 and *L* is the size of the simulation cell. Multiple trial displacements are proposed until a trial position that does not result in steric clash for the entire molecule is generated. The move is then attempted. As with the slithering snake move, each monomer in the molecule that is translated will have an orientational bias. Accordingly, the Rosenbluth factors are calculated as in equations (14) and (15). The translational move results in large displacements for single chains and correctly biases the system for efficient sampling of configurations with alternate interaction patterns.

Translational moves can also be applied to clusters of molecules. A connected cluster refers to a collection of unique chains connected via rotational interactions. A proposed move only results in a translation, and the move is readily accepted if there is no steric clash. Since no new physical bonds are created at the proposed location, the cluster remains invariant and the move is accepted. Naively this move might seem unnecessary as this move simply moves clusters around. However, once a physical bond has formed between two molecules, it is highly unlikely for any of the non-cluster translation moves to move the centers-of-masses of clusters closer together. Cluster moves are essential in that they mimic the Ostwald ripening process whereby large droplets consume smaller ones (67). Without cluster moves, ripening is quenched because a single large droplet cannot form, especially at lower simulation temperatures.

### Identifying phase boundaries using measures for density inhomogeneities

In order to detect the onset of phase separation, we can calculate excess chemical potentials using the Widom particle insertion method (68) and equalize these chemical potentials across distinct phases. This process requires *a priori* knowledge of the densities of both phases. An efficient variant of this approach, based on fast Fourier transforms, was recently developed and deployed by Qin and Zhou (69). They demonstrated their method for calculation of liquid-liquid coexistence curves for a patchy colloid model of *γ*II-cyrstallin. Given that LASSI simulations are lattice-based and that we do not have *a priori* knowledge of the density within the dense phase that forms upon phase separation, we instead rely on properties of pair distribution functions that help us diagnose the onset of phase separation and compute phase boundaries. Pair distribution functions are helpful because phase separation is the result of the system partitioning into phases of different densities. The pair distribution, which is a reduced-dimension partition function, serves as a rigorous thermodynamic and structural measure of local density and inhomogeneities of density. To first order, the density fluctuations are quantified by averaging over all orientations. Accordingly, the pair distribution function can be converted to a radial distribution function that allows us to probe local densities and local structural organization of molecules around one another. However, the normalization of the pair distribution function requires some caution. The system contains polymer molecules and using a prior distribution that assumes an ideal gas of the chain monomers to normalize the pair distribution function is problematic because it does not accurately capture the effects of non-idealities due to chain connectivity. We leverage the efficient sampling of polymer fluids in LASSI and obtain suitable prior distributions by simulating the system of interest in the absence of sticker-sticker interactions.

The pair distribution function *P*^(2)^(*r*) quantifies the equilibrium distribution of distances between chain monomers, where *r* is the inter-monomer distance. If 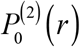 denotes the prior pair distribution function calculated from simulations where the inter-sticker interactions are ignored (see **Fig 7a**), then the normalized radial distribution function is written as:

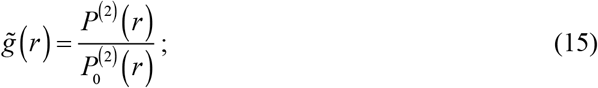

**Fig 7.**
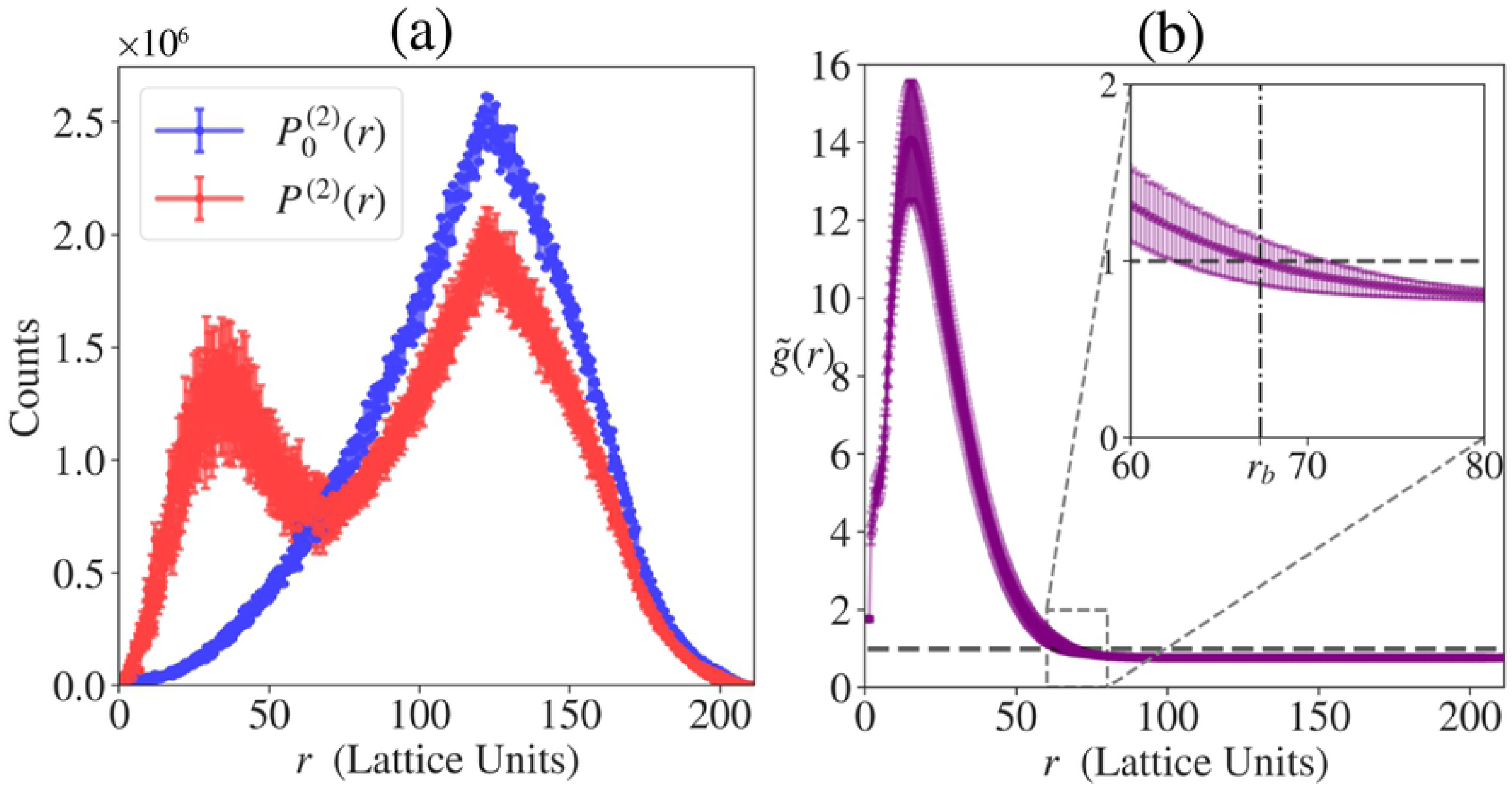
Distribution functions used for calculation of density inhomogeneity. The data shown are obtained from 5 independent A_*n*_-B_*n*_ simulations with total protein concentration *c* = 6.89×10^−5^ voxels^−1^ and reduced temperature *T*\* = 0.383. Error bars indicate standard deviations. (a) Pair distribution functions *P*^(2)^(*r*) and *P*_0_^(2)^(*r*), where the former is from the interacting system and the latter from the non-interacting system with chain connectivity (*prior pair distribution function*). Note that *P*^(2)^(*r*) shows two peaks, the first of which indicates dense phase formation. (b) Radial distribution function 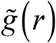. This captures the droplet formation by a sharp and broad peak in the beginning. The inset shows *r*_b_ where 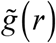 intersects the line corresponding to 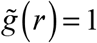 line, delineating between the dense and solution phases. The global density inhomogeneity measure, ξ, is obtained by integration of absolute deviation of 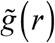 from 1.

The function 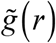 is a direct measure of the local density of the protein of interest. Since LASSI uses periodic boundary conditions, the maximal inter-monomer distance is 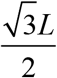. Given this normalized 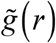, we note that if the system has short-range ordering like a canonical liquid, the radial distribution function will oscillate around unity but eventually approach one as *r* → ∞. Conversely, if the system undergoes a density transition, 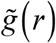 will have two distinct spatial regimes (**Fig 7b**): for small *r*, 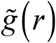 will be characterized by a tall and broad peak such that 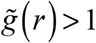 until 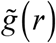 intersects the 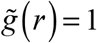 line; this region corresponds to the dense phase and we shall denote the value of *r* at this intersection to be *r* = *r*_b_. For *r* > *r*_b_, 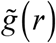 will be between 0 and 1, and for lattices that are large enough to avoid finite size artifacts, 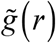 will converge to a value lower than one that corresponds to the dilute phase region. Furthermore, 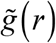 can be used to estimate the densities within the dense and dilute phases.

To quantify the global density inhomogeneity we introduce a simple measure, ξ, which is calculated as follows:

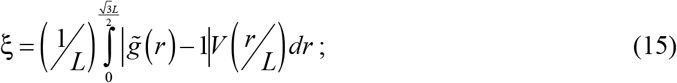

In equation (17), *V*(*r*/*Lx*), the volume element for the normalized radial distance *x*=*r*/*L* defined as:

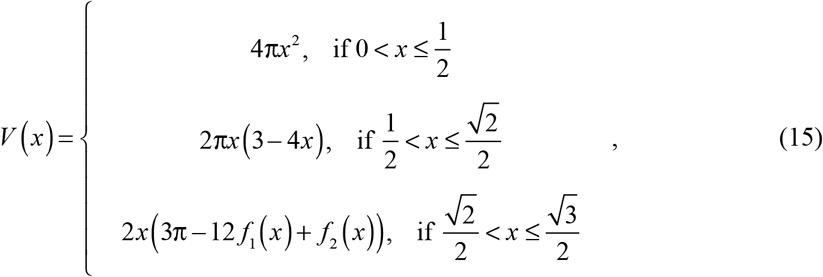

and

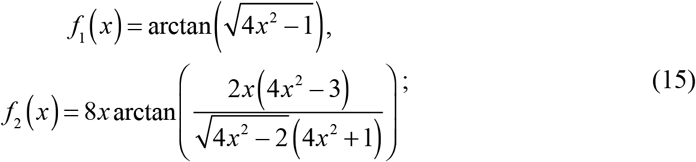

If ξ ≈ 0, the global density inhomogeneity in the system is small and this will be characteristic of a single homogeneous phase dominating the simulation volume. As ξ increases beyond zero, the system accommodates density inhomogeneities. Currently, to draw the phase diagrams we use a cutoff value of ξ = 0.025, which is universal to all systems, to delineate between a homogeneous system and one that has phase separated.

### Estimating the percolation transition line that delineates percolated and nonpercolated networks

Associative polymers form networks characterized by physical crosslinks among stickers. Accordingly, we use the concept of a *cluster, viz*., a collection of unique chains connected via rotational interactions, to define the extent of percolation. In polymer melt simulations, the extent of percolation, known as the *gel fraction* in the polymer literature, is defined as the fraction of polymers participating in a percolating network that spans the simulation box in at least one direction (70). More generally, we can use the fraction of polymers that make up the single largest cluster to quantify the onset of percolation and the changes to the extent of networking beyond the percolation threshold (71). A molecular network cannot percolate the whole simulation cell when dilute and dense phases coexist. Accordingly, we choose the second definition for the order parameter that describes the percolation transition, and we denote this as ϕ_c_ (29).

Semenov and Rubinstein demonstrated that a percolation transition is not a thermodynamic phase transition, but is instead purely a connectivity transition (45). This implies that the identification of the percolation threshold is not achievable using a *bona fide* order parameter but instead relies on a suitable topological description. Here, we employ the midpoint of the ϕ_c_ vs. concentration curve to assess the onset of percolation and the percolation line is obtained as the locus of points in the phase diagram for which ϕ_c_ = 0.5. In a system where finite size artifacts are minimized, the percolation transition is sharp having either a hyperbolic or sigmoidal shape as a function of concentration. Accordingly, the location of the percolation line will be relatively robust to the choice one makes for the percolation threshold.

## Results

We demonstrate the use of LASSI by applying it to study two archetypal systems that are known to undergo phase separation (24, 31, 33). The systems are instantiations of linear and branched multivalent protein systems, respectively. The simulation results obtained for linear multivalent proteins illustrate how phase diagrams are generated when protein concentration (at a fixed stoichiometry) and temperature are the independent variables. In the second example that includes a branched multivalent protein and a linear peptide, the temperature is fixed, and the concentrations of the individual components are varied. The simulation parameters used in both simulations are summarized in **Table 1**. For each system, we conducted 5 independent simulations, each of which consists of 5×10^8^ MC steps after 5×10^6^ equilibration steps. The data were taken over the last half of the trajectories at a frequency of 5×10^5^ steps.

**Table 1.** Simulation parameters for system description.

|  | Linear multivalent system<br>(see **Figure 8c**) | Branched multivalent system<br>(see **Figure 12b**) |
| --- | --- | --- |
| Bead notations | A/B: stickers<br>N: neutral spacers | A/B: stickers<br>N: neutral hub spacers |
| Number of stickers $s_i$ | $s_A = 5, s_B = 5$ | $s_A = 10, s_B = 5$ |
| Linker length $l_{ij}$<br>(in lattice units) | $l_{AN} = 1, l_{BN} = 1, l_{NN} = 4$ | $l_{NA} = 1, l_{AA} = 3, l_{BB} = 3$ |
| Position-dependent energy<br>$E_{\text{pos}}(\mathbf{r}_1, \mathbf{r}_2)$ | ∞, if $\mathbf{r}_1 = \mathbf{r}_2$<br>0, otherwise | ∞, if $\mathbf{r}_1 = \mathbf{r}_2$<br>0, otherwise |
| Pairwise interaction energy $\varepsilon_{ij}$<br>(in temperature units) | $\varepsilon_{AB} = -3, \varepsilon_{ii} = 0, \varepsilon_{iN} = 0$ | $\varepsilon_{AB} = -3, \varepsilon_{ii} = 0, \varepsilon_{iN} = 0$ |

### FUS as an example of a linear multivalent protein

Wang *et al*. (24) recently uncovered the molecular grammar that contributes to the driving forces for phase separation of the FUS family proteins. They showed that, to first order, 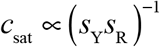, where *c*_sat_ is the measured saturation concentration of the FUS family proteins and *s*_Y_ and *s*_R_ are the number of tyrosine (Tyr) and arginine (Arg) residues, respectively. In FUS and other proteins under investigation, the Tyr residues are located primarily within the N-terminal disordered prion-like domain (PLD), whereas the Arg residues are located primarily within the partially disordered C-terminal RNA binding domain (RBD).

Wang *et al*. showed that Tyr and Arg residues are the stickers in the FUS family proteins. Accordingly, the zeroth-order stickers and spacers representation used to model FUS in LASSI comprises of two parts: An N-terminal mimic of the PLD encompassing Tyr residues as stickers and a C-terminal mimic of the RBD that encompasses Arg residues as stickers. Wang *et al*. also measured *c*_sat_ for a 1:1 mixture of independent PLDs and RBDs interacting in *trans*. The *c*_sat_ for this system is approximately twice that of the *c*_sat_ for full-length FUS. Given the block copolymeric architecture of FUS, we denote the PLD and RBD as A_n_ and B_n_, respectively for A and B-blocks of valence n. The model system of PLDs and RBDs interacting in *trans* is denoted as A_n_+B_n_ (**Fig 8a**), whereas the system mimicking full-length FUS where the stickers can interact in *cis* and in *trans* is denoted as A_n_-B_n_ (**Fig 8b**).

**Fig 8.**
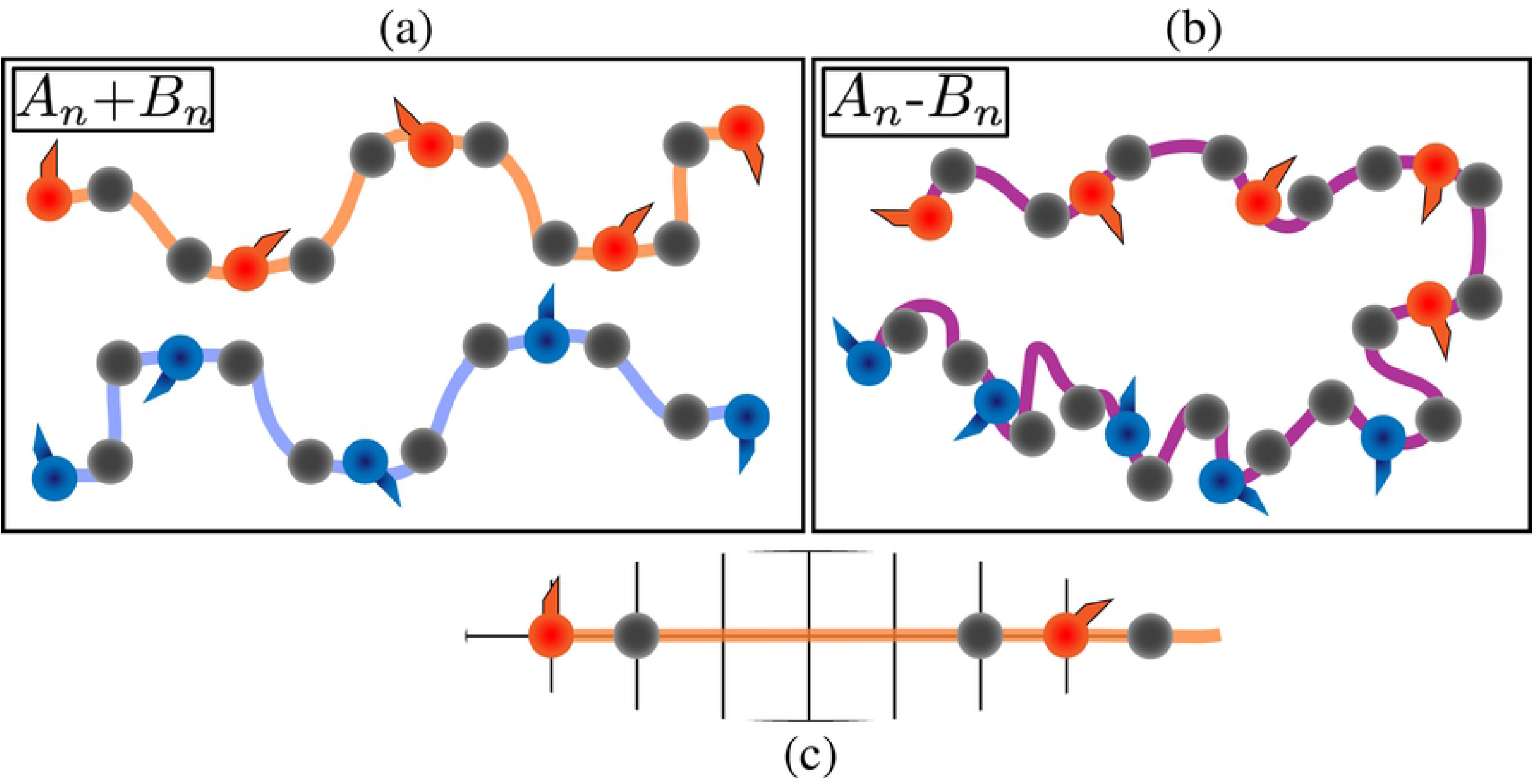
Architecture of the linear multivalent systems. (a, b) Cartoons to depict the A_*n*_+B_*n*_ and A_*n*_-B_*n*_ systems, respectively. Different colors of beads denote different species of stickers. Note that A_*n*_-B_*n*_ can be simply considered as A_*n*_+B_*n*_ where the two different sections of the proteins are joined together. (c) Linker lengths involved in the architecture (see also **Table 1**). Each sticker has a neighboring spacer bead that is 1 lattice site apart whereas the neighboring spacer beads are 4 lattice sites apart. This means that consecutive stickers are 6 lattice sites apart and also that the linkers connecting the two have a positive effective solvation volume.

Within An and Bn blocks, spacers provide a uniform spacing of six lattice sites between stickers along the chain. We model spacers using a hybrid approach whereby a neutral spacer monomer is attached to each sticker with spacing of a single lattice site (**Fig 8c**). This choice was made to provide a distinction between A_n_-B_n_ and A_n_+B_n_. Accordingly, the relative concentration of neutral beads will be higher in A_n_-B_n_ when compared to A_n_+B_n_. This allows us to incorporate linker-mediated differences between the driving forces for phase separation for A_n_-B_n_ vs. A_n_+B_n_

#### Computed phase diagrams are consistent with experiments

**Fig 9** shows phase diagrams for the A_n_+B_n_ and A_n_-B_n_ systems calculated using LASSI. In panels (a) and (b), the phase diagram is projected onto a plane, where the ordinate quantifies the reduced temperatures *T*^*^ calculated as 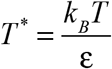 where ε is the effective energy of pairwise interactions between stickers from the A_n_ and B_n_ blocks. Panel (a) shows results for the A_n_+B_n_ system. The bulk concentration in the A_n_+B_n_ system is quantified along the abscissa as 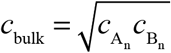 where *c*_A_n__ and *c*_B_n__ are the bulk concentrations of A_n_ and B_n_, respectively. Panel (b) shows the phase diagram for the A_n_-B_n_ system where the abscissa represents the bulk concentration of this system.

**Fig 9.**
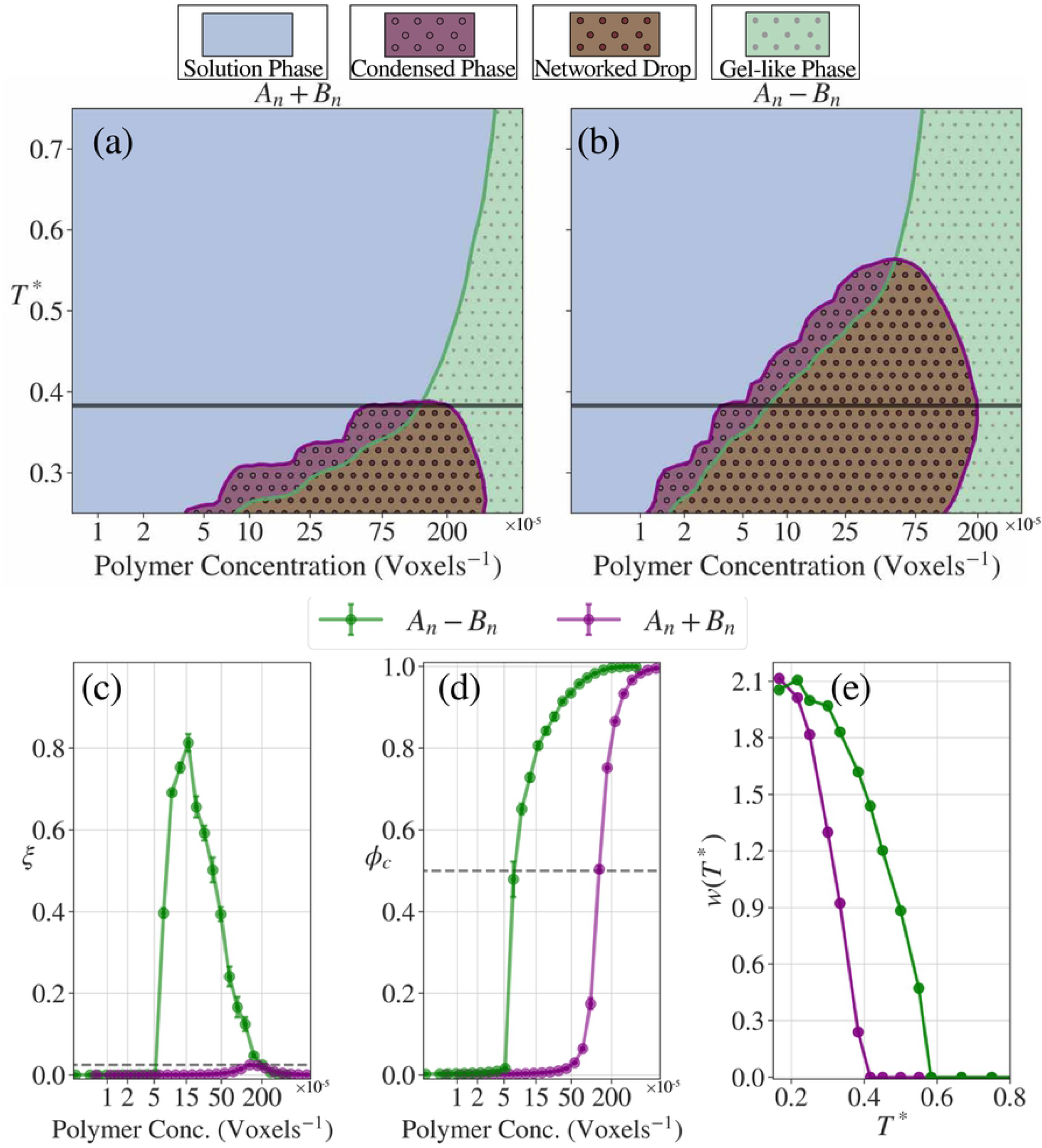
Phase behavior of the linear multivalent systems. (a, b) Phase diagrams for the A_*n*_+B_*n*_ and A_*n*_-B_*n*_ systems, respectively. The purple line is a 2-dimensional linear interpolation for ξ = 0.025, and the area encapsulated by the purple line are where the systems have large density inhomogeneities and are thus considered to be phase separated. The green line is a 2-dimensional linear interpolation for ϕ_c_ = 0.5 and thus is the proxy for the percolation line. (c, d) ξ and ϕ_c_ curves as a function of concentrations at *T*\* = 0.383 (solid lines in (a) and (b)). (e) Width of the two-phase regime, *w*(*T*\*), as a function of the reduced temperature. Not only does the A_*n*_-B_*n*_ system have a higher critical temperature (*T*\* ~ 0.6 vs. *T*\* ~ 0.4), but also has a wider two-phase regime than the A_*n*_+B_*n*_ system.

Experiments show that the driving forces for phase separation are roughly twice as large for the full-length FUS compared to the system comprising of a 1:1 mixture of PLDs and RBDs (24). This feature is recapitulated in LASSI simulations. For example, the width of the two-phase regime is larger for the A_n_-B_n_ system compared to the A_n_+B_n_ system for all values of *T*^*^ as shown in panel (c) of **Fig 9**. The critical temperature is higher for the A_n_-B_n_ vs. A_n_+B_n_ system (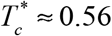 vs. 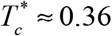, respectively). The valence of stickers is the main determinant of the concentration at the critical point whereas the interactions mediated by spacers determine the density inhomogeneities and hence the critical temperature. The impact of the longer chain length and increased valence of stickers per chain is also evident from the percolation threshold, which is crossed at approximately two-fold lower bulk protein concentrations for the A_n_-B_n_ system when compared to the A_n_+B_n_ system, across all the simulation temperatures. Differences between the two systems are also evident in the degree of cooperativity of phase separation and the percolation transition as shown in panels (d) and (e) of **Fig 9**.

For each system, the intersection of the percolation threshold line with the two-phase regime shows that the dense phase predominantly forms a percolated droplet (panels (a) and (b) in **Fig 9**). Accordingly, for associative polymers, the condensed phase sans percolation within the droplet is a metastable phase. Unlike homopolymers, which are polymers made up entirely of stickers or spacers, associative polymers comprise of a mixture of stickers and spacers. Stickers provide the driving forces for networking and the spacers ensure that these driving forces lead to condensation. The importance of the sticker-driven percolation is evidenced in the persistence of percolated networks for both systems at high values of *T*^*^.

The observation that dense phases form percolated droplets has several implications: (1) on timescales that are concordant with or smaller than the average lifetime of physical crosslinks between stickers, the condensate will have elastic properties; this will be replaced by viscous behavior on timescales that are longer than the average lifetime of physical crosslinks (72); (2) condensates will have an intrinsic tendency for viscoelasticity (73) and long-lived crosslinks will cause hardening behavior that is observed in many systems (1, 6, 7, 11, 22, 24, 39–41); (3) the extent of crosslinking above the percolation threshold will change continuously with concentration (16, 29), and this will govern the overall structure, mechanics, and porosity of condensates; (4) reactions within condensates are likely to be constrained by the network topology formed as a result of inter-sticker interactions (74); these constraints can create a variety of interesting kinetic signatures for reactions (75), including temporal memories as has been demonstrated recently for a system that undergoes thermoresponsive phase behavior (76). The results obtained from LASSI simulations provide a richer assessment of the overall phase behavior and further analysis of these phase transitions, across a multitude of similar systems should lead to the development of coherent theoretical explanations for the comparative driving forces for phase transitions of linear multivalent proteins.

#### Move set frequencies and diagnostics of converged simulations

We used results from simulations of the linear multivalent protein system to assess the design of LASSI. The frequencies of the different move sets for simulations of the linear multivalent protein system are summarized in **Table 2**. Considerations that go into the design of move sets include the achievement of converged equilibrium distributions, with maximal computational efficiency, for each bulk concentration. Diagnostics from short simulations are often useful to optimize the move set frequencies especially if multiple short trials are performed using very different starting configurations.

**Table 2.** Move frequencies according to their types. They are normalized to the sum of all frequencies used in each simulation.

|  | Linear multivalent system | Branched multivalent system |
| --- | --- | --- |
| Cluster translation move | 1 | 1 |
| Chain translation move | 10 | 10 |
| Rotation move | 100 | 100 |
| Local move | 250 | 250 |
| Reptation move | 0 | 50 |
| Double pivot move | 50 | 10 |

The structure of each move set also serves as a guide for selecting an optimal set of frequencies. This leads to a set of heuristics that are as follows: (i) the cluster move picks a random chain from the system, performs a networking analysis on that chain, and then proposes a displacement of the cluster. As the cluster size grows it is more likely that a randomly picked chain will be part of the largest cluster which itself will probably result in a steric clash after the proposed move. Therefore, the frequency of the cluster move should be low, if not the lowest, in the entire set. (ii) The translational move picks a random chain from the system for translation; as the size of the largest cluster increases it becomes less likely for a proposed translation move to be accepted. However, unlike the cluster move the translation move is rotationally biased and thus results in new interactions being formed. Hence, translational moves enable single-molecule to cluster-surface interactions. Therefore, this move should be proposed more often than the cluster move, although not as often as rotational or local moves. (iii) The rotation move is computationally inexpensive and it enables the switching of physical bonds and should thus be proposed fairly frequently. (iv) Similarly, local moves and slithering snake moves are also rotationally biased, and they help with the local rearrangements of physical bonds. Local moves are the primary route to enable local conformational changes, and to enable local physical bond rearrangements. Therefore, local moves should be proposed most frequently. The slithering snake move is particularly effective because it allows for large local physical bond rearrangements in dense configurations. Thus, this move should also be proposed frequently, less so than local moves but more so than translation moves. Note that in a system where some molecules are non-linear or have heterogeneous linker lengths, the frequency would need to be higher since the move is rejected if an incorrect molecule is picked at random. (v) The double pivot move allows for large-scale changes to conformations within dense configurations and accordingly, this move should be proposed more frequently than both cluster and translation moves. One can track the acceptance ratios of each move over a very rough initial sweep across the relevant system parameters. Moves that are always rejected do not enable any changes in configuration and only add computational costs. Therefore, the frequency for that particular move should be lowered. This is especially the case for the cluster move in high-density systems.

**Fig 10** shows the concentration dependence of acceptance ratios for each of the move types, diagnosed for simulations of the A_n_+B_n_ and A_n_-B_n_ systems. The acceptance ratios show similar qualitative trends for both systems, even though there are clear quantitative differences between the systems. The move with the highest acceptance ratio in the dense regime is the double pivot move, signifying that the systems are transitioning into a pure polymer melt where the proteins become increasingly entangled. The second most accepted move is the local move; extrapolating from the higher concentrations it is expected that the acceptance of local moves should also become small and that the double pivot move is the most dominant interacting mode, since even to move one monomer, multiple monomers from multiple chains would need to be moved. Both systems have similar qualitative trends for the translation move where we see a transition from being accepted at low concentrations to not being accepted at higher concentrations. Since the proteins in the A_n_-B_n_ system are twice long as the A_n_+B_n_ system, the absolute acceptance ratio of the translation move in always lower in the A_n_-B_n_ system.

**Fig 10.**
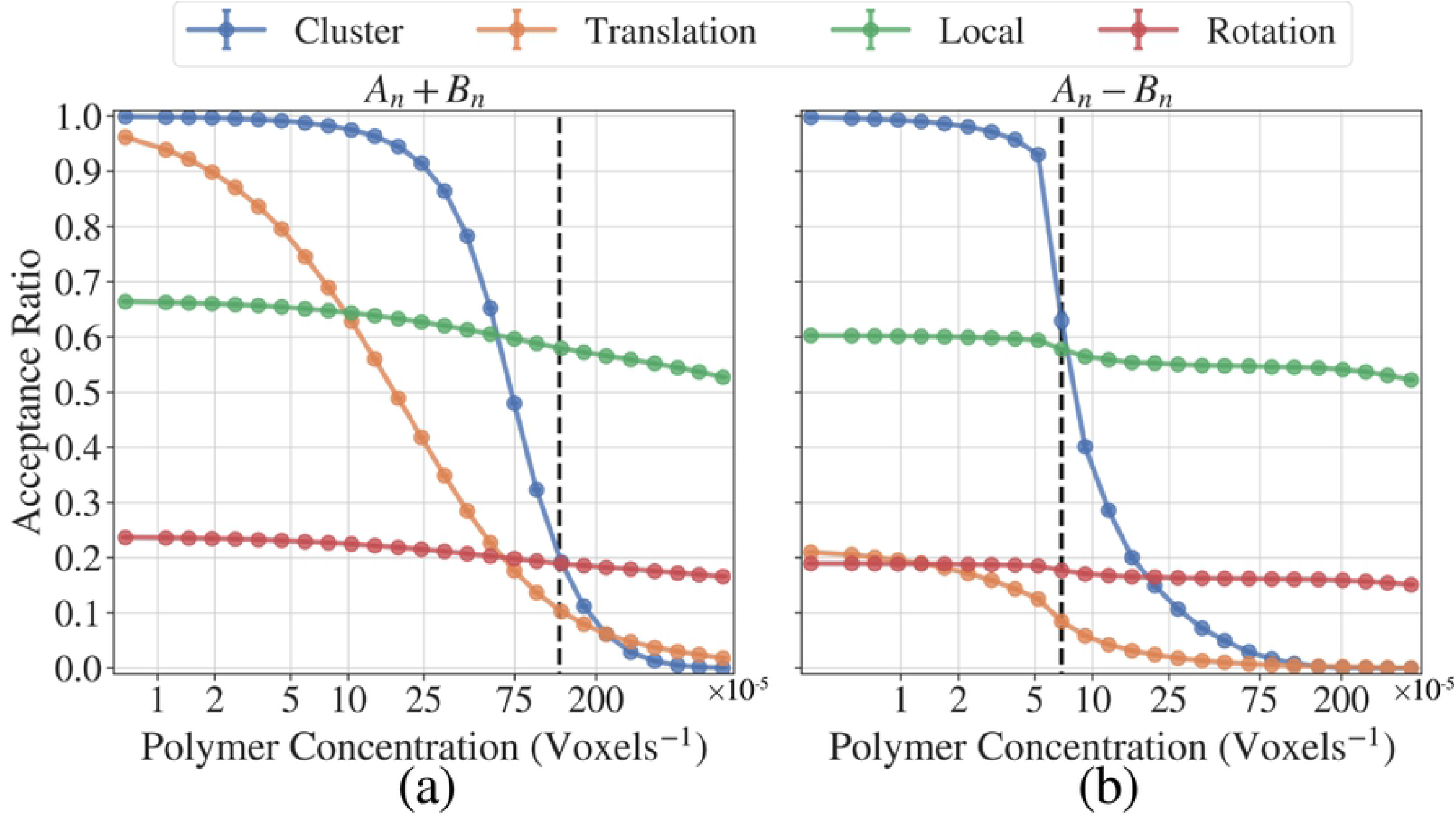
Move acceptance ratios. Curves with different colors indicate acceptance ratios of different types of moves. The dashed lines show the saturation concentrations. The data are obtained from simulations with *T*\* = 0.383.

Cluster moves have high acceptance ratios in the dilute regime whereas the acceptance ratio nearly vanishes as the concentration increases. This is intuitive since the likelihood of steric clashes increases with a decrease in available volume and this is coupled to the simultaneous increase in the fraction of molecules in the largest cluster. We note here that the cluster moves have the most dramatic change in acceptance ratios from values near 1 to values near 0. This behavior is expected since the cluster move is the primary mechanism by which the condensates coarsen. However, the apparent inefficiency of cluster moves in dense configurations cannot be used as a rationale to quench such moves. In fact, as shown in **Fig 11**, phase separation is suppressed if cluster moves are not part of the move set. This highlights the importance of cluster moves for generating *bona fide* phase separation within finite-sized systems and has a bearing on the collective coordinate over which density inhomogeneities are created and grown in systems that undergo phase separation.

**Fig 11.**
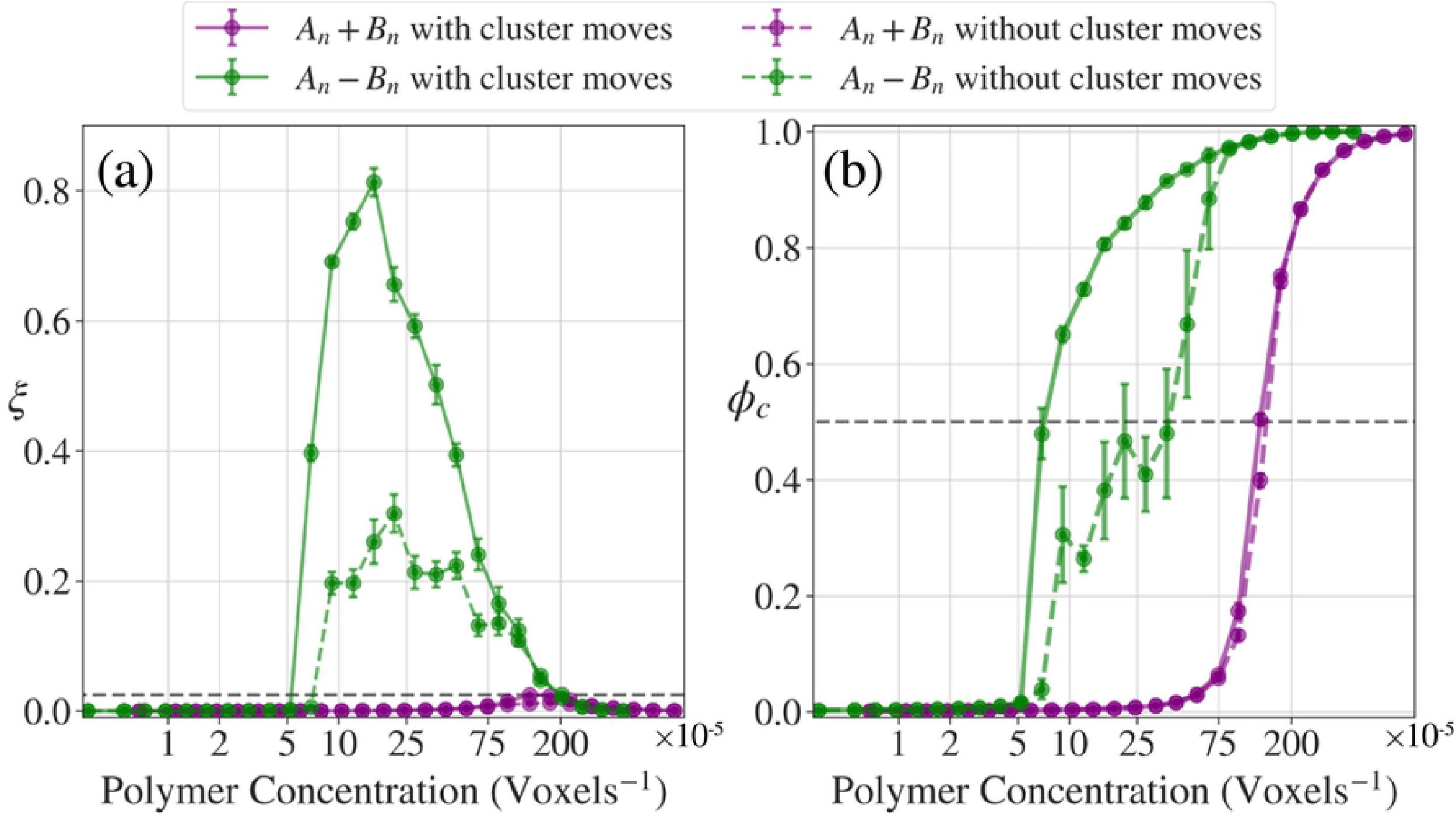
Importance of cluster translation moves. (a)ξ and (b) ϕ_c_ curves for the A_*n*_+B_*n*_ (purple) and A_*n*_-B_*n*_ systems (green) at *T*\* = 0.383. The sold lines are identical with the curves in panels (c) and (d) of **Figure 9**. The dotted lines show the simulation results under the same system conditions but the frequency for cluster translation moves is set to zero. Not only do the systems phase separate and percolate at higher saturation concentrations, but also we can see that both percolation and separation are suppressed highly. Furthermore, errors are generally higher, due to the systems being highly dependent on the initial conditions of the system.

### Mixtures of N130 and the rpL5 peptide as an example of a branched multivalent protein system

LASSI can also be deployed to study the phase behavior of branched multivalent proteins. Examples of branched multivalent proteins are molecules that form symmetric, stable oligomers such as nucleophosmin 1 (NPM1) and synthetic systems such as the corelets designed by Bracha et al. (77). NPM1 is a key molecule within the granular component (GC) of the nucleolus (78). Three coexisting layers define the nucleolus and the GC is the outermost layer. Within the GC, NPM1 appears to form facsimiles of percolated droplets in complex ribosomal proteins with Arg-rich motifs (17, 30). A minimalist system that mimics the phase behavior of the GC comprises of a truncated version of NPM1, referred to as N130, and an Arg-rich peptide, designated as rpL5 (31–33). Both NPM1 and N130 form symmetric pentamers in the presence of Arg-rich peptides (79). The pentamer formed by the association of folded domains serves as a scaffold for displaying disordered C-terminal tails that are defined by two distinct acidic tracts. The system also features an N-terminal disordered region with a well-defined acidic motif.

In the LASSI representation, N130 pentamers with disordered tails are modeled using a five-armed structure. This approach follows the computational strategy of Feric *et al*. (30), which was based on the fact that pentamers do not dissociate under conditions where NPM1 / N130 undergo phase separation. Each arm comprises of two sticker sites to mimic the presence of the A1 and A2 acidic tracts within the disordered tails of NPM1 / N130. Therefore, each N130 pentamer displays a total of ten sticker sites. The spacers between each A1 tract and the N130 core as well as between each pair of A1 and A2 tracts on a disordered tail are phantom tethers, which means that their effective solvation volumes (29) are set to zero. Each rpL5 peptide has two sticker sites corresponding to the two Arg-rich motifs along the sequence. Schematic representations of the minimalist coarse-grained architecture used for N130 and rpL5 are shown in **Fig 12**.

**Fig 12.**
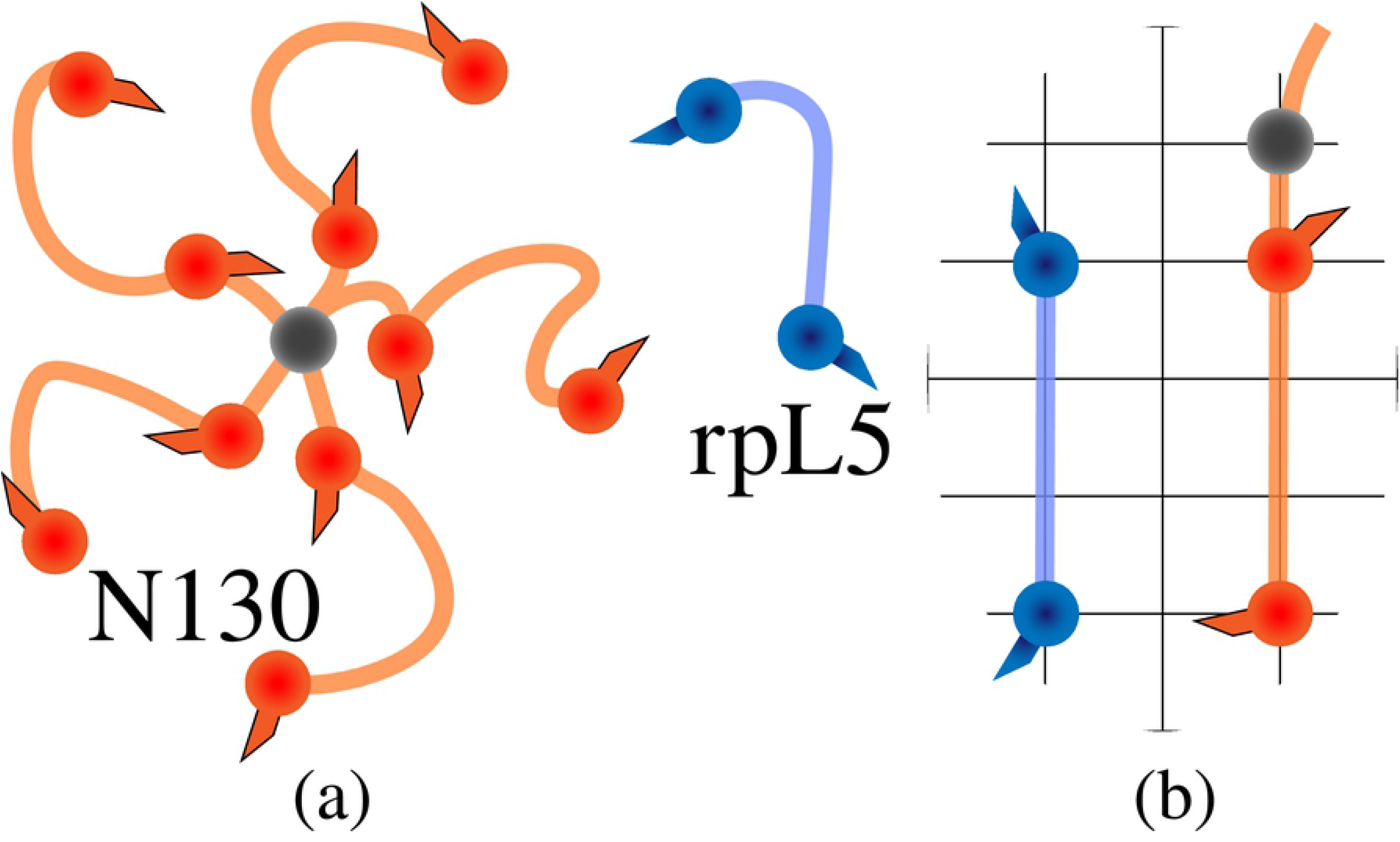
Architecture of an archetypal branched multivalent system. (a) Schematic to depict the overall architecture. The pentamer with 10 orange stickers represents the N130 molecule where the gray central oligomerization domain is modeled as a neutral spacer monomer, and the rpL5 peptide is modeled as a linear molecule with 2 blue stickers. (b) Relevant length scales for the architecture (see also **Table 1**). For the rpL5 molecule a linker length of 3 was chosen between the two stickers, and for the N130 molecule the first sticker (modeling the A1 tract) is 1 lattice site away from the hub spacer whereas the second sticker (modeling the A2 tract) is 3 lattice sites away from the first sticker.

#### Computed phase diagrams and their geometric shapes

Given that the phase behavior of the N130 + rpL5 system is driven by heterotypic interactions involving the A1 / A2 tracts from the N130 tails and the Arg-motifs from rpL5, we constructed phase diagrams by keeping the simulation temperature fixed and varied the concentrations of N130 and rpL5 molecules. Projections of phase diagrams onto planes defined by N130 concentration along the abscissa and rpL5 concentration along the ordinate are shown in panel (a) of **Fig 13**. The general shape of the phase diagram is comparable with that of the experimentally determined phase diagram (33), even though direct comparison is not straightforward because the scarcity of experimental data points does not yield a full phase diagram. Therefore, instead of direct comparison with experiment, we provide a detailed discussion on the shapes of the percolation line and the phase boundary.

**Fig 13.**
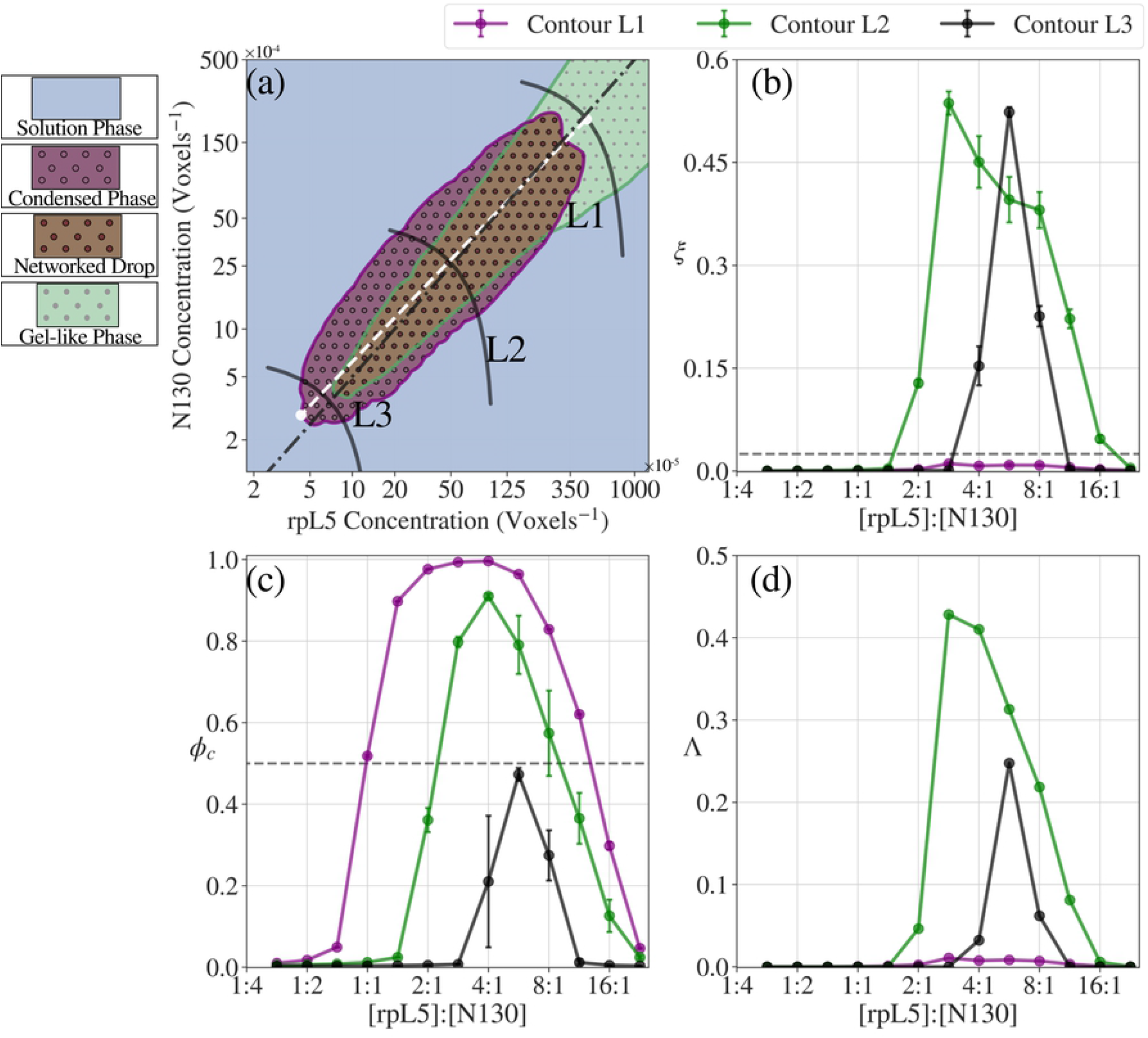
Phase behaviors of the branched multivalent systems for *T*\* = 0.25. (a) Full phase diagram, where the purple line denotes the proxy for the binodal and the green line is the proxy for the percolation line (see also the caption for **Figure 9**). The phase-separated region has an elliptical shape and we have a closed loop, which demonstrates re-entrant phase behavior, whereas the percolation line has a conical shape extending into much higher densities. The solid black lines denote contours of constant total concentration where L1 is the lowest concentration and L3 is the highest concentration. Note that both axes are represented in the log scale. (b, c) ξ and ϕ_c_ curves as a function of relative stoichiometric ratio of N130 and rpL5 along the constant-concentration contours. (d) Plot of Λ vs. the apparent stoichiometry along lines L1, L2, and L3.

The percolation line, constructed as a function of the concentrations of two multivalent molecules, has an overall parabolic shape. This feature may be explained as follows: Let *f*_1_ and *f*_2_ denote the fractions of N130 and rpL5 molecules that are bound in a network; *s*_1_ and *s*_2_ will denote the corresponding valence of stickers on N130 and rpL5, respectively (for the current implementation of the N130 + rpL5 system, *s*_1_ = 10 and *s*_2_ = 2). The percolation threshold is crossed when *f*_1_*f*_2_(*s*_1_–1)(*s*_2_–1) > 1. If we keep (*s*_1_–1)(*s*_2_–1) constant, the threshold concentration for percolation will be governed by the product *f*_1_*f*_2_. Accordingly, if there is a large excess of N130 (component 1) compared to rpL5 (component 2), then *f*_1_ ← 0 and *f*_2_ ← 1, and the system does not undergo a percolation transition. In this scenario, every rpL5 molecule is crosslinked to two sticker sites from one or two N130 molecules. However, since the relative stoichiometry favors N130 molecules, there is a vast excess of un-crosslinked N130 molecules and the network cannot grow. Percolation is also inhibited when the converse situation arises, *i.e*., when there is a large excess of component 2 with respect to component 1. Accordingly, the percolation line takes on a parabolic form in the plane defined by the concentrations of N130 and rpL5.

While the percolation line is parabolic, the phase boundary, defined by the density transition, is elliptical thus forming a closed loop in the plane defined by the concentrations *c*_1_ and *c*_2_ of N130 and rpL5, respectively. In associative polymers, the phase behavior is governed by the affinity between stickers, the valence of stickers, and the effective solvation volumes of spacers (28, 29). For fixed *c*_1_ that intersects the two-phase regime an increase in *c*_2_ will show evidence of two saturation concentrations for component 2 (rpL5). The lower saturation concentration corresponds to the first density transition governed by *c*_2_ approaching *c*_1_. Beyond the second saturation concentration, there is a growing excess of rpL5 molecules and not enough N130 molecules to drive the density transition via inter-sticker interactions. Similar reasoning applies to describe the reentrant behavior that will result by keeping *c*_2_ fixed at a value that intersects the two-phase regime and increasing *c*_1_. The density transition will have a closed loop structure because the differences between the saturation concentrations for the two components decrease with increasing values of *c*_bulk_. Therefore, beyond an apparent threshold value of *c*_bulk_, the system will exit into the homogeneous one-phase regime. A portion of this one-phase regime will be defined by percolation without phase separation as shown in panel (a) of **Fig 13**.

Taken together, the parabolic percolation lines and elliptic forms for two-phase regimes define conic sections that highlight the reentrant phase behavior whereby fixing the concentration of component 1 and increasing the concentration of the second species can lead to phase separation and percolation at a low threshold concentration of component 2 and exit into the one-phase, non-percolated regime beyond a second higher threshold concentration for component 2. This type of reentrant phase behavior has been reported for a model protein + RNA system (80). It is worth emphasizing, however, that reentrant phase behavior will be a generic attribute of multicomponent systems comprising of associative polymers. It is not unique to protein / RNA systems or systems governed by electrostatic interactions. These findings, which emerge naturally from LASSI simulations, should enable the development of a general theory for the shapes of multidimensional phase diagrams and predictions for novel ways to regulate phase transitions within cells.

#### Apparent stoichiometric ratios can be different from effective stoichiometric ratios

Stoichiometry of scaffold molecules is another key parameter that determines the functions of biomolecular condensates formed by multicomponent systems (81). The apparent stoichiometric ratio is calculated as the ratio of the concentrations of stickers of types *s*_1_ and *s*_2_ for N130 and rpL5, respectively such that 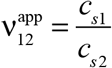. However, the effective stoichiometric ratio 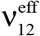 can be different from 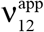 if excluded volume effects modulate the effective concentration of stickers. We fit an ellipse to the two-phase boundary and determined the major axis of this ellipse. The effective stoichiometric ratio should be unity along the major axis. As shown in panel (a) of **Fig 13**, the major axis deviates from the line along which 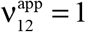. Therefore, 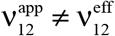 and angle between the major axis and the line along which 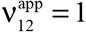 quantifies the impact of excluded volume on changes to effective concentrations of stickers that in turn modifies the stoichiometric ratios. This calculation demonstrates the utility of having access to the full phase diagram, which is made possible by LASSI.

The synergy between stoichiometry and phase behavior can be analyzed by quantifying the order parameter ξ and the topological parameter ϕ_c_ as a function of apparent stoichiometry for fixed bulk concentration. Along each gray line in panel (a) of **Fig 13** the total concentration defined as *c*_bulk_ = (*c*_1_*c*_2_)^½^ is fixed, although the stoichiometries will vary. The mean values of *c*_bulk_ along L1, L2, and L3 are 2.09×10^−2^ (voxel^−1^), 2.46×10^−3^ (voxel^−1^), and 3.33×10^−4^ (voxel^−1^), respectively and the value of 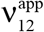 ranges from 0.36 to 22.62 along each of L1, L2, and L3. Panels (b) and (c) in **Fig 13** show the variation of ξ and ϕ_c_ as 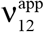 increases along L1, L2, and L3, respectively. Along L1, the value of ξ is essentially zero irrespective of stoichiometry because L1 lies above the second threshold point and is outside the two-phase regime. However, a system spanning percolated network forms for stoichiometries in the range 1.2 ≤ ν_12_ ≤ 13 along L1. This is because the concentrations of both components are well above the percolation threshold along L1 thus ensuring that stickers readily find one another even without a density transition. In direct contrast, along L3, we observe phase separation, characterized by values of ξ > 0.025 for a range of stoichiometries, but none of these support the formation of a percolated droplet (ϕ_c_ < 0.5 for all stoichiometries). Along L2, we observe phase separation for stoichiometries in the range 1.15 ≤ ν_12_ ≤ 16 and percolation for stoichiometries in the range 2.14 ≤ ν_12_ ≤ 11.3 such that phase separation enables the formation of a percolated droplet.

In panel (d) of **Fig 13**, we introduce a new structural parameter Λ, which we define as a convolution of ξ and ϕ_c_ such that Λ = ξ⊗ϕ_c_. Here, the convolution is calculated as a logical AND gate, which becomes a simple product. The parameter Λ quantifies the convolution of the density and network transition and quantifies the extent to which the phase separation and percolation are coupled as the apparent stoichiometry is varied for a fixed bulk concentration. The profile of Λ is reminiscent of the profile measured by Case *et al*. (81) for the dwell time of signaling molecules as a function of stoichiometric ratios that govern the formation of condensates at membranes. This suggests that dwell times, which are experimentally accessible parameters, might in fact be proxies for a more fundamental quantity, viz., the structural features of the condensates as measured by the convolution between phase separation and percolation and the extent of network formation within the condensate.

The key finding is that the combination of the bulk concentration and stoichiometric ratio (as opposed to stoichiometry alone) will determine the quench depth into the two-phase regime. This in turn determines whether a system-spanning network forms without phase separation or if phase separation enables the formation of a droplet-spanning network. The structure of condensates and the overall phase behavior cannot be fully described in terms of *c*_bulk_ or ν_12_ alone. Instead, this requires the consideration of both parameters jointly and relative to the saturation concentrations as well as the percolation threshold. This is important because the extent of crosslinking and the time scales associated with crosslinks will determine the material properties of the condensate. This in turn should contribute to parameters such as the dwell times of clients within condensates (81) as well as the temporal and spatial control of the exchange clients between condensates and the surrounding milieu.

## Discussion

In this work, we have built on the established connection between multivalent proteins and associative polymers (44, 45, 74) with their stickers-and-spacers architecture (15, 17, 24, 28, 29, 40, 43, 47) to develop and deploy LASSI, a lattice-based open source computational engine that enables the simulation of system-specific phase diagrams of single and multi-component systems. The foundations of LASSI derive from the formalism of the bond fluctuation model (57, 58, 82) and what we report here is a formalization of recent *ad hoc* deployments LASSI. Also, we demonstrated the application of LASSI to archetypal linear and branched multivalent proteins. These are pseudo one- and two-component systems inasmuch as the sites on lattices not occupied by protein modules are in fact solvent sites. This allows us to capture the effects of mixing entropies, through the excluded volume of vacant sites. Further, the pairwise interaction strengths between protein sites are effective interactions that represent the three-way interplay among protein-protein, protein-solvent, and solvent-solvent interactions.

We demonstrate how canonical ensemble Monte Carlo simulations with appropriately designed move sets and analysis of order parameters derived from the distribution functions allow us to calculate full phase diagrams as a function of protein concentration and interaction strengths. These phase diagrams help us capture the interplay between sticker driven and spacer mediated phase separation and percolation, a distinctive feature of associative polymers (29, 45, 72). LASSI allows us to query the impact of overall and intrinsic valence of stickers, interaction strengths between stickers, the spatial ranges of these interactions, the effective solvation volumes and lengths of spacers, and protein concentrations. These titrations generate multidimensional phase diagrams. Here, we have shown how insights can be gleaned by projecting multidimensional phase diagrams onto the two-dimensional space. Going forward, it would be helpful to generate mathematical and geometrical descriptions of phase behavior in higher than two dimensions, especially since all biomolecular condensates have hundreds of distinct molecular components. The calculation of pair and higher order distribution functions should afford multiscale descriptions of the structural organization of molecular components within condensates. The acceptance ratios associated with different move sets and the length scales spanned by distinct move sets open the door to analyzing the dynamics of phase separation, percolation and the interplay between the two. It should also allow us to explore the dynamics of molecular exchange and the dynamical evolution of interaction stoichiometries within condensates. Another major direction of future application for LASSI is to uncover the determinants of compositional specificity of condensates (1, 12).

The choice of a lattice-based approach for coarse-graining and modeling phase behavior of multivalent proteins is guided by the established advantages of lattice models (83) for polymeric systems and the disadvantages of off-lattice coarse-grained models. To titrate across the full range of volume fractions, one needs to balance considerations of finite size artifacts – which requires large numbers of molecules – with large simulation volumes – which makes it difficult to observe density fluctuations that can grow into density inhomogeneities. Stylized, slab-like geometries need to be incorporated into off-lattice simulations to enable the calculation of full phase diagrams (84). Such approaches raise concerns about using *O*(10^2^) and confined volumes to generate phase diagrams. Lattices help us work around these problems because the conformational space is discretized, the calculation of interaction potentials can be made to be very efficient through the use of look up tables, and a better balance between limiting finite size artifacts and the need for large simulation volumes can be achieved because of the optimal scaling of computational cost with system size. Additionally, lattice-based simulations are readily generalized to incorporate directional interactions, increased intrinsic and overall valence, modeling the interplay between density and networking transitions, and the incorporation of multiple components.

The approaches deployed in LASSI have been applied to model a variety of multicomponent systems, including mimics of RNA molecules (24, 28-30, 43, 53). These approaches allow for hybrid representations and can be generalized to systems of arbitrary complexity. What is required is the development of approaches that enable systematic coarse-graining and adaptation of machine learning based methods to parameterize interaction potentials (85). Engineering LASSI to be interoperable to cell-based modeling suites (86) will also allow for larger scale deployment of the overall framework.

## Acknowledgments

This work was supported by grants from the US National Science Foundation (MCB-1614766), the Human Frontier Science Program (RGP0034/2017), the US National Institutes of Health (5R01NS056114), and the St. Jude Children's Research Hospital through the research collaborative on membraneless organelles. The funders had no role in study design, data collection and analysis, decision to publish, or preparation of the manuscript. We thank Tyler Harmon for his original contributions, helpful discussions, and critical insights. We are also grateful to Simon Alberti, Martin Fossat, Alex Holehouse, Anthony Hyman, Tanja Mittag, Michael Rosen, Kiersten Ruff, and Andrea Soranno for many helpful discussions. LASSI is designed to be an open source engine that will be available via http://pappulab.wustl.edu that will provide a hyperlink to a suitable code hosting, reviewing, and distribution site.

## References

1. Banani SF, Lee HO, Hyman AA, Rosen MK. Biomolecular condensates: organizers of cellular biochemistry. Nature Reviews Molecular Cell Biology. 2017;18(5):285–98.

2. Shin Y, Brangwynne CP. Liquid phase condensation in cell physiology and disease. Science. 2017;357(6357):eaaf4382.

3. Lamond AI, Spector DL. Nuclear speckles: a model for nuclear organelles. Nature Reviews in Molecular and Cell Biology. 2003;4(8):605–12.

4. Mintz PJ, Spector DL. Compartmentalization of RNA processing factors within nuclear speckles. Journal of structural biology. 2000;129(2-3):241–51.

5. Brangwynne CP, Eckmann CR, Courson DS, Rybarska A, Hoege C, Gharakhani J, et al. Germline P granules are liquid droplets that localize by controlled dissolution/condensation. Science. 2009;324(5935):1729–32.

6. Saha S, Weber CA, Nousch M, Adame-Arana O, Hoege C, Hein MY, et al. Polar positioning of phase-separated liquid compartments in cells regulated by an mRNA competition mechanism. Cell. 2016;166(6):1572–84.e16.

7. Patel A, Lee HO, Jawerth L, Maharana S, Jahnel M, Hein MY, et al. A liquid-to-solid phase transition of the ALS protein FUS accelerated by disease mutation. Cell. 2015;162(5):1066–77.

8. Su X, Ditlev JA, Hui E, Xing W, Banjade S, Okrut J, et al. Phase separation of signaling molecules promotes T cell receptor signal transduction. Science. 2016;352(6285):595–9.

9. Banjade S, Rosen MK. Phase transitions of multivalent proteins can promote clustering of membrane receptors. eLife. 2014;3:e04123.

10. Zeng M, Chen X, Guan D, Xu J, Wu H, Tong P, et al. Reconstituted Postsynaptic Density as a Molecular Platform for Understanding Synapse Formation and Plasticity. Cell. 2018;174(5):1172–87.e16.

11. Hyman AA, Weber CA, Jülicher F. Liquid-liquid phase separation in biology. Annu Rev Cell Dev Biol. 2014;30:39–58.

12. Banani SF, Rice AM, Peeples WB, Lin Y, Jain S, Parker R, et al. Compositional Control of Phase-Separated Cellular Bodies. Cell. 2016;166(3):651–63.

13. Wu H, Fuxreiter M. The Structure and Dynamics of Higher-Order Assemblies: Amyloids, Signalosomes, and Granules. Cell. 2016;165(5):1055–66.

14. Falkenberg CV, Carson JH, Blinov ML. Multivalent Molecules as Modulators of RNA Granule Size and Composition. Biophysical Journal. 2017;113(2):235–45.

15. Posey AE, Holehouse AS, Pappu RV. Phase Separation of Intrinsically Disordered Proteins. In: Rhoades E, editor. Methods in Enzymology. 611: Academic Press; 2018. p. 1–30.

16. Li P, Banjade S, Cheng H-C, Kim S, Chen B, Guo L, et al. Phase transitions in the assembly of multivalent signalling proteins. Nature. 2012;483(7389):336–40.

17. Wei MT, Elbaum-Garfinkle S, Holehouse AS, Chen CC, Feric M, Arnold CB, et al. Phase behaviour of disordered proteins underlying low density and high permeability of liquid organelles. Nature Chemistry. 2017;9(11):1118–25.

18. Zhang H, Elbaum-Garfinkle S, Langdon EM, Taylor N, Occhipinti P, Bridges AA, et al. RNA controls PolyQ protein phase transitions. Molecular Cell. 2015;60(2):220–30.

19. Elbaum-Garfinkle S, Kim Y, Szczepaniak K, Chen CC-H, Eckmann CR, Myong S, et al. The disordered P granule protein LAF-1 drives phase separation into droplets with tunable viscosity and dynamics. Proceedings of the National Academy of Sciences USA. 2015;112(23):7189–94.

20. Molliex A, Temirov J, Lee J, Coughlin M, Kanagaraj AP, Kim HJ, et al. Phase Separation by Low Complexity Domains Promotes Stress Granule Assembly and Drives Pathological Fibrillization. Cell. 2015;163(1):123–33.

21. Posey AE, Ruff KM, Harmon TS, Crick SL, Li A, Diamond MI, et al. Profilin reduces aggregation and phase separation of huntingtin N-terminal fragments by preferentially binding to soluble monomers and oligomers. Journal of Biological Chemistry. 2018;293(10):3734–46.

22. Mateju D, Franzmann TM, Patel A, Kopach A, Boczek EE, Maharana S, et al. An aberrant phase transition of stress granules triggered by misfolded protein and prevented by chaperone function. The EMBO Journal. 2017:e201695957.

23. Rai AK, Chen J-X, Selbach M, Pelkmans L. Kinase-controlled phase transition of membraneless organelles in mitosis. Nature. 2018;559(7713):211–6.

24. Wang J, Choi J-M, Holehouse AS, Lee HO, Zhang X, Jahnel M, et al. A Molecular Grammar Governing the Driving Forces for Phase Separation of Prion-like RNA Binding Proteins. Cell. 2018;174(3):688–99.e16.

25. Lin Y-H, Brady J P, Forman-Kay J D, Chan HS. Charge pattern matching as a ‘fuzzy’ mode of molecular recognition for the functional phase separations of intrinsically disordered proteins. New Journal of Physics. 2017;19(11):115003.

26. Pak CW, Kosno M, Holehouse AS, Padrick SB, Mittal A, Ali R, et al. Sequence determinants of intracellular phase separation by complex coacervation of a disordered protein. Mol Cell. 2016;63(1):72–85.

27. Brangwynne CP, Tompa P, Pappu RV. Polymer physics of intracellular phase transitions. Nature Physics. 2015;11(11):899–904.

28. Harmon TS, Holehouse AS, Pappu RV. Differential solvation of intrinsically disordered linkers drives the formation of spatially organized droplets in ternary systems of linear multivalent proteins. New Journal of Physics. 2018;20(4):045002.

29. Harmon TS, Holehouse AS, Rosen MK, Pappu RV. Intrinsically disordered linkers determine the interplay between phase separation and gelation in multivalent proteins. eLife. 2017;6:e30294.

30. Feric M, Vaidya N, Harmon TS, Mitrea DM, Zhu L, Richardson TM, et al. Coexisting liquid phases underlie nucleolar subcompartments. Cell. 2016;165(7):1686–97.

31. Mitrea DM, Cika JA, Stanley CB, Nourse A, Onuchic PL, Banerjee PR, et al. Self-interaction of NPM1 modulates multiple mechanisms of liquid-liquid phase separation. Nature Communications. 2018;9(1):842.

32. Ferrolino MC, Mitrea DM, Michael JR, Kriwacki RW. Compositional adaptability in NPM1-SURF6 scaffolding networks enabled by dynamic switching of phase separation mechanisms. Nature Communications. 2018;9(1):5064.

33. Mitrea DM, Cika JA, Guy CS, Ban D, Banerjee PR, Stanley CB, et al. Nucleophosmin integrates within the nucleolus via multi-modal interactions with proteins displaying R-rich linear motifs and rRNA. eLife. 2016;5:e13571.

34. Brady JP, Farber PJ, Sekhar A, Lin Y-H, Huang R, Bah A, et al. Structural and hydrodynamic properties of an intrinsically disordered region of a germ cell-specific protein on phase separation. Proceedings of the National Academy of Sciences. 2017;114(39):E8194–E203.

35. Vernon RM, Chong PA, Tsang B, Kim TH, Bah A, Farber P, et al. Pi-Pi contacts are an overlooked protein feature relevant to phase separation. eLife. 2018;7:e31486.

36. Nott Timothy J, Petsalaki E, Farber P, Jervis D, Fussner E, Plochowietz A, et al. Phase Transition of a Disordered Nuage Protein Generates Environmentally Responsive Membraneless Organelles. Molecular Cell. 2015;57(5):936–47.

37. Lin Y, Currie SL, Rosen MK. Intrinsically disordered sequences enable modulation of protein phase separation through distributed tyrosine motifs. Journal of Biological Chemistry. 2017.

38. Boeynaems S, Bogaert E, Kovacs D, Konijnenberg A, Timmerman E, Volkov A, et al. Phase Separation of C9orf72 Dipeptide Repeats Perturbs Stress Granule Dynamics. Mol Cell. 2017;65(6):1044–55.e5.

39. Woodruff JB, Ferreira Gomes B, Widlund PO, Mahamid J, Honigmann A, Hyman AA. The Centrosome Is a Selective Condensate that Nucleates Microtubules by Concentrating Tubulin. Cell. 2017;169(6):1066–77.e10.

40. Franzmann TM, Jahnel M, Pozniakovsky A, Mahamid J, Holehouse AS, Nüske E, et al. Phase separation of a yeast prion protein promotes cellular fitness. Science. 2018;359(6371):eaao5654.

41. Maharana S, Wang J, Papadopoulos DK, Richter D, Pozniakovsky A, Poser I, et al. RNA buffers the phase separation behavior of prion-like RNA binding proteins. Science. 2018;360(6391):918–21.

42. Langdon EM, Qiu Y, Ghanbari Niaki A, McLaughlin GA, Weidmann CA, Gerbich TM, et al. mRNA structure determines specificity of a polyQ-driven phase separation. Science. 2018;360(6391):922–7.

43. Boeynaems S, Holehouse AS, Weinhardt V, Kovacs D, Van Lindt J, Larabell C, et al. Spontaneous driving forces give rise to protein-RNA condensates with coexisting phases and complex material properties. Proceedings of the National Academy of Sciences. 2019:https://doi.org/10.1073/pnas.1821038116.

44. Rubinstein M, Dobrynin AV. Solutions of Associative Polymers. Trends in Polymer Science. 1997;5(6):181–6.

45. Semenov AN, Rubinstein M. Thermoreversible Gelation in Solutions of Associative Polymers. 1. Statics. Macromolecules. 1998;31(4):1373–85.

46. Halabi N, Rivoire O, Leibler S, Ranganathan R. Protein Sectors: Evolutionary Units of Three-Dimensional Structure. Cell. 2009;138(4):774–86.

47. Holehouse AS, Pappu RV. Functional Implications of Intracellular Phase Transitions. Biochemistry. 2018;57(17):2415–23.

48. Rubinstein M, Colby RH. Polymer Physics. New York: Oxford University Press; 2003 2003.

49. Stockmayer WH. Theory of Molecular Size Distribution and Gel Formation in Branched - Chain Polymers. The Journal of Chemical Physics. 1943;11(2):45–55.

50. Flory PJ. Molecular Size Distribution in Three Dimensional Polymers. I. Gelation1. Journal of the American Chemical Society. 1941;63(11):3083–90.

51. Flory PJ. Constitution of Three-dimensional Polymers and the Theory of Gelation. The Journal of Physical Chemistry. 1942;46(1):132–40.

52. Dias CS, Araújo NAM, Telo da Gama MM. Dynamics of network fluids. Advances in Colloid and Interface Science. 2017;247:258–63.

53. Fei J, Jadaliha M, Harmon TS, Li IT, Hua B, Hao Q, et al. Quantitative analysis of multilayer organization of proteins and RNA in nuclear speckles at super resolution. Journal of Cell Science. 2017;130(24):4180–92.

54. Harmon TS, Holehouse AS, Pappu RV. To Mix, or To Demix, That Is the Question. Biophysical Journal. 2017;112(4):565–7.

55. Freeman Rosenzweig ES, Xu B, Kuhn Cuellar L, Martinez-Sanchez A, Schaffer M, Strauss M, et al. The Eukaryotic CO<sub>2</sub>-Concentrating Organelle Is Liquidlike and Exhibits Dynamic Reorganization. Cell. 2017;171(1):148–62.e19.

56. Bolhuis P, Frenkel D. Numerical study of the phase diagram of a mixture of spherical and rodlike colloids. The Journal of Chemical Physics. 1994;101(11):9869–75.

57. Shaffer JS. Effects of chain topology on polymer dynamics: Bulk melts. The Journal of Chemical Physics. 1994;101(5):4205–13.

58. Carmesin I, Kremer K. The bond fluctuation method: a new effective algorithm for the dynamics of polymers in all spatial dimensions. Macromolecules. 1988;21(9):2819–23.

59. Metropolis N, Rosenbluth AW, Rosenbluth MN, Teller AH, Teller E. Equation of State Calculations by Fast Computing Machines. The Journal of Chemical Physics. 1953;21(6):1087–92.

60. Siepmann JI, Frenkel D. Configurational bias Monte Carlo: a new sampling scheme for flexible chains. Molecular Physics. 1992;75(1):59–70.

61. Rosenbluth MN, Rosenbluth AW. Monte Carlo Calculation of the Average Extension of Molecular Chains. The Journal of Chemical Physics. 1955;23(2):356–9.

62. Mańka A, Nowicki W, Nowicka G. Monte Carlo simulations of a polymer chain conformation. The effectiveness of local moves algorithms and estimation of entropy. Journal of Molecular Modeling. 2013;19(9):3659–70.

63. Qin Y, Liu H-L, Hu Y. Dynamic Monte Carlo Simulation of Polymers: Cooperative Move Algorithm. Molecular Simulation. 2003;29(10-11):649–54.

64. Gennes PGd. Reptation of a Polymer Chain in the Presence of Fixed Obstacles. The Journal of Chemical Physics. 1971;55(2):572–9.

65. Sheng Y-J, Wang M-C. Statics and dynamics of a single polymer chain confined in a tube. The Journal of Chemical Physics. 2001;114(10):4724–9.

66. Binder K, Paul W. Monte Carlo simulations of polymer dynamics: Recent advances. Journal of Polymer Science Part B: Polymer Physics. 1997;35(1):1–31.

67. Vladimirova N, Malagoli A, Mauri R. Diffusion-driven phase separation of deeply quenched mixtures. Physical Review E. 1998;58(6):7691–9.

68. Widom B. Some Topics in the Theory of Fluids. The Journal of Chemical Physics. 1963;39(11):2808–12.

69. Qin S, Zhou H-X. Fast Method for Computing Chemical Potentials and Liquid–Liquid Phase Equilibria of Macromolecular Solutions. The Journal of Physical Chemistry B. 2016;120(33):8164–74.

70. Rottereau M, Gimel JC, Nicolai T, Durand D. Monte Carlo simulation of particle aggregation and gelation: I. Growth, structure and size distribution of the clusters. The European Physical Journal E. 2004;15(2):133–40.

71. Mikes J, Dusek K. Simulation of polymer network formation by the Monte Carlo method. Macromolecules. 1982;15(1):93–9.

72. Koyama T, Tanaka H. Volume-shrinking kinetics of transient gels as a consequence of dynamic interplay between phase separation and mechanical relaxation. Physical Review E. 2018;98(6):062617.

73. Shayegan M, Tahvildari R, Kisley L, Metera K, Michnick SW, Leslie SR. Probing inhomogeneous diffusion in the microenvironments of phase-separated polymers under confinement. bioRxiv. 2018:402230.

74. Rubinstein M, Semenov AN. Thermoreversible Gelation in Solutions of Associating Polymers. 2. Linear Dynamics. Macromolecules. 1998;31(4):1386–97.

75. Kopelman R. Fractal Reaction Kinetics. Science. 1988;241(4873):1620–6.

76. Roberts S, Harmon TS, Schaal JL, Miao V, Li K, Hunt A, et al. Injectable tissue integrating networks from recombinant polypeptides with tunable order. Nature Materials. 2018;17(12):1154–63.

77. Bracha D, Walls MT, Wei M-T, Zhu L, Kurian M, Avalos JL, et al. Mapping Local and Global Liquid Phase Behavior in Living Cells Using Photo-Oligomerizable Seeds. Cell. 2018;175(6):1467–80.e13.

78. Sirri V, Urcuqui-Inchima S, Roussel P, Hernandez-Verdun D. Nucleolus: the fascinating nuclear body. Histochemistry and Cell Biology. 2008;129(1):13–31.

79. Mitrea DM, Grace CR, Buljan M, Yun M-K, Pytel NJ, Satumba J, et al. Structural polymorphism in the N-terminal oligomerization domain of NPM1. Proceedings of the National Academy of Sciences. 2014;111(12):4466–71.

80. Banerjee PR, Milin AN, Moosa MM, Onuchic PL, Deniz AA. Reentrant Phase Transition Drives Dynamic Substructure Formation in Ribonucleoprotein Droplets. Angewandte Chemie International Edition. 2017;56(38):11354–9.

81. Case LB, Zhang X, Ditlev JA, Rosen MK. Stoichiometry controls activity of phase-separated clusters of actin signaling proteins. Science. 2019;363(6431):1093–7.

82. Subramanian G, Shanbhag S. On the relationship between two popular lattice models for polymer melts. The Journal of Chemical Physics. 2008;129(14):144904.

83. Kremer K, Binder K. Monte Carlo simulation of lattice models for macromolecules. Computer Physics Reports. 1988;7(6):259–310.

84. Dignon GL, Zheng W, Kim YC, Best RB, Mittal J. Sequence determinants of protein phase behavior from a coarse-grained model. PLOS Computational Biology. 2018;14(1):e1005941.

85. Ruff KM, Harmon TS, Pappu RV. CAMELOT: A machine learning approach for coarsegrained simulations of aggregation of block-copolymeric protein sequences. The Journal of Chemical Physics. 2015;143(24):243123.

86. Resasco DC, Gao F, Morgan F, Novak IL, Schaff JC, Slepchenko BM. Virtual Cell: computational tools for modeling in cell biology. Wiley Interdisciplinary Reviews: Systems Biology and Medicine. 2012;4(2):129–40.

